# B cells expressing authentic naive human VRC01-class BCRs can be primed and recruited to germinal centers in multiple independent mouse models

**DOI:** 10.1101/2020.02.24.963629

**Authors:** Deli Huang, Robert K. Abbott, Colin Havenar-Daughton, Patrick D. Skog, Rita Al-Kolla, Bettina Groschel, Tanya R. Blane, Sergey Menis, Jenny Tuyet Tran, Theresa C. Thinnes, Sabrina A. Volpi, Mark Pintea, James E Voss, Nicole Phelps, Ryan Tingle, Alberto R. Rodriguez, Greg Martin, Sergey Kupryianov, William R. Schief, David Nemazee, Shane Crotty

**Author notes:** These authors contributed equally (Deli Huang, Robert K. Abbott).

## Abstract

Animal models of human antigen-specific B cell receptors (BCR) generally depend on “inferred germline” sequences, and thus their relationship to authentic naive human B cell BCR sequences and affinities is unclear. Here, BCR sequences from authentic naive human VRC01-class B cells from healthy human donors were selected for the generation of three new BCR knock-in mice. The BCRs span the physiological range of affinities found in humans, and use three different light chains (VK3-20, VK1-5, and VK1-33) found among subclasses of naive human VRC01-class B cells and HIV broadly neutralizing antibodies (bnAbs). The germline-targeting HIV immunogen eOD-GT8 60mer is currently in clinical trial as a candidate bnAb vaccine priming immunogen. To attempt to model human immune responses to the eOD-GT8 60mer, we tested each authentic naive human VRC01-class BCR mouse model under rare human physiological B cell precursor frequency conditions. B cells with high (HuGL18^HL^) or medium (HuGL17^HL^) affinity BCRs were primed, recruited to germinal centers, accrued substantial somatic hypermutation, and formed memory B cells. Precursor frequency and affinity interdependently influenced responses. Taken together, these experiments utilizing authentic naive human VRC01-class BCRs validate a central tenet of germline-targeting vaccine design and extend the overall concept of the reverse vaccinology approach to vaccine development.

## INTRODUCTION

Vaccinology has largely been an empirical science over the past century. While this approach has produced many successful vaccines (*1, 2*), the lack of vaccines to pathogens like HIV has suffered from our lack of knowledge of the fundamental biology of how B cells compete in vivo following immunization (*3*). Nearly all vaccines are thought to work by induction of protective antibody responses (*4*). There is increasing optimism that effective vaccine strategies for HIV might be found. Recent efforts have focused on recurrent broadly neutralizing antibody (bnAb) specificities, such as the VRC01 class, that might be elicited by vaccines (*5*). The concept of reverse vaccinology (aka, reverse vaccinology 2.0) involves engineering proteins as immunogens based on knowledge of bnAbs, or protective mAbs more broadly (*6, 7*). Succinctly put, reverse vaccinology defines antibody-to-vaccine strategies of vaccine design. Germline-targeting vaccine design is a vaccine concept based on designing a priming immunogen specifically capable of binding B cell receptors (BCRs) of “germline” precursors of bnAbs, based on BCR sequence relationships, and thereby eliciting related B cell responses in vivo (*8*). One apparent barrier to eliciting bnAbs is that the inferred germline (iGL) BCR precursors of most bnAbs have negligible affinity for HIV Env (*5*), and so are subdominant. BnAb precursors are also predicted to be quite rare (*9*). Competing B cells reactive to non-conserved epitopes tend to dominate the response (*3*) and fail to provide protection, as evidenced during natural infection by the ubiquity of strain-specific antibodies and the rarity of bnAbs (*10*).

In response to these challenges, germline-targeting immunogens have been engineered. First generation germline-targeting immunogens have been developed based on inferred unmutated ancestor, i.e. iGL, bnAb sequences (*8, 11–18*). Development of candidate vaccines in that manner has two challenges that relate to BCR sequences and the human B cell repertoire. First, authentic gl-bnAb BCR sequences are almost always difficult to predict (*14, 19–21*). Second, the protein engineering was frequently focused on binding to a single (or small set of) bnAb-precursor sequence match(es) that may not sufficiently relate to sequences normally found in the human B cell repertoire. Germline-targeting requires the ability to target BCRs of precursors of the desired bnAb, specifically BCRs actually found in the human naive B cell repertoire. While knock-in mouse models have provided substantial value in understanding the in vivo challenges that exist in maturing B cells towards broad neutralization (*11, 12, 22–27*), it is unclear if these models are capable of revealing all of the hurdles that authentic VRC01-class B cells will face with germline-encoded CDR3s. Second generation germline-targeting immunogens have incorporated broader sets of precursors of representative bnAbs and BCR sequences identified from NGS sequencing of HIV-negative donors predicted to be related to bnAb-precursors (*28, 29*). VRC01-class germline-targeting immunogens have progressed furthest in development by these criteria. eOD-GT8 60mer is a germline-targeting HIV Env engineered outer domain nanoparticle that has now entered human clinical trials (*30*).

VRC01-class bnAbs can neutralize upwards of 98% of HIV viral isolates. The VRC01 class is special in several respects (*31, 32*). Broad and potent owing to its favorable angle of approach and because it mimics interactions of CD4 (engaging the CD4 binding site of HIV Env (CD4bs)), the VRC01 class uses a single VH gene, VH1-2, along with a critical short 5 amino acid (aa)-long L-CDR3 (typically QQYEF). To develop neutralizing breadth, VCR01-class Abs also require acquisition of a small L-CDR1 deletion or other accommodation to avoid a clash with the Env glycan at N276 (*9, 19, 33*). VRC01-class B cells undergo 19-40% aa somatic hypermutation (SHM) from germline to develop into bnAbs in HIV^+^ individuals (*32–36*). Germinal centers (GCs) are the anatomical site in which antigen-activated B cells undergo SHM, clonal competition, and Darwinian selection by T follicular helper (T_FH_) cells for survival (*37–40*). These rounds of mutation and selection produce mutated high affinity antibodies in a process known as affinity maturation (*41, 42*). VRC01-class B cells will have to compete successfully in these processes in response to immunization if a germline-targeting VRC01-class bnAb vaccine is to succeed.

One way to rigorously assess the quality of a germline-targeting immunogen is to determine if it can identify bnAb precursor naive B cells in humans, via direct binding. To date this has only been accomplished for two germline-targeting immunogen designs (*22, 28, 29, 43, 44*). We discovered that VRC01-class B cells (i.e. B cells that contain the VH1-2*02 [or VH1- 2*04], heavy chain (HC) allele paired with a light chain (LC) with a 5 aa L-CDR3) do exist in the naive B cell repertoire of most humans, at a frequency of 1 in 300,000 B cells, and those B cells can be identified by their binding to eOD-GT8 (*43*). VRC01-class eOD-GT8-bound human naive B cells could be grouped into subclasses, based on LC V gene. eOD-GT8-binding VRC01-class naive human B cells of different subclasses are predicted to differ in their ability to develop into bnAbs, based on the ease with which they accommodate the HIV Env N276 glycan (*43*). VK1-5^+^ or VK1-33^+^ VRC01-class B cells can likely do so by point mutations within L-CDR1, whereas VK3-20^+^ VRC01-class B cells appear to require L-CDR1 deletions or longer H-CDR3s (*19, 32, 45*). Further investigation of the VRC01-class human naive B cell repertoire revealed the BCRs vary widely in affinity for eOD-GT8, H-CDR3 sequence, and H-CDR3 length (*43*). Importantly, it is not yet known whether subsets of these eOD-GT8-binding VRC01-class naive human B cells can respond in vivo, and it is not yet known whether they differ in their abilities to compete in vivo.

Mice do not possess a VH1-2 gene homolog, and thus cannot make VRC01-class Abs, necessitating the analysis in various types of knock-in mice. eOD-GT8 60mer has shown promise as a priming immunogen in mice carrying an inferred germline BCR of the original VRC01 bnAb (VRC01^gH^ or VRC01^gHL^) (*12, 27, 44, 46*) or the 3BNC60 bnAb HC (*11*), or in recombining HC knock-in mice carrying the VH1-2 gene (*22, 44*). Given that VRC01-class precursors are rare in humans, we developed a VRC01^gHL^ mouse B cell transfer strategy to better model human physiological precursor frequencies and affinities (*27*). Precursor frequency and antigen affinity were both shown to be important factors in determining B cell competitive success in GCs (*27*). The VRC01^gHL^ model relied on an iGL bnAb sequence, which has limitations, both because the sequence was used in the original germline-targeting immunogen design process and because the L-CDR3 and H-CDR3 sequences were relatively unchanged from the VRC01 bnAb because these regions are not germline encoded.

Here we created three different BCR knock-in H/L mouse models carrying authentic VRC01-class BCRs identified from isolated human naive B cells. We found that VRC01-class B cells from all three BCR knock-in H/L mouse models were able to be primed and participate in GCs when adoptively transferred at high precursor frequencies. Upon transfer into congenic recipients to achieve a human-like physiologically reasonable VRC01-class precursor frequency of ∼10^-6^, high affinity and intermediate affinity VRC01-class naive B cells responded in vivo, participated in GCs, and accrued SHMs, including VRC01-class mutations. Taken together, these data validate a key tenet of the germline-targeting approach to vaccine design. We have shown for the first time that authentic human BCR expressing B cells can be successfully primed in vivo to HIV germline-targeting antigens when starting from rare precursor frequencies matched to the human physiological range. These data suggest that eOD-GT8 60mer, and germline-targeting immunogens with similar properties, are likely to be successful in human clinical trials.

## RESULTS

### Generation of three BCR knock-in mouse models with authentic antigen-specific human naive B cell BCR specificity: HuGL16, HuGL17, and HuGL18

We generated mouse models carrying three different VRC01-class BCRs identical to those from human naive B cell donors. These are termed “HuGL” mice due to their expression of authentic antigen-specific “<u>hu</u>man <u>g</u>ermline” naive B cell BCR specificities. All use VH1-2*02, JH4*02, and LCs with 5 aa-long CDRL3 C<u>QQY(E/D)X</u> motif but vary in H-CDR3 length, affinity for eOD-GT8, and VK usage (**Fig 1A**, **Table S1**). HuGL16 (K_D_ 18.5 μM) was chosen for its use of VK1-33, which is capable of Env N276 glycan-accommodating glycine mutations in L-CDR1 rather than deletions. HuGL17 (Kd 1.3 μM) uses VK1-5, which should similarly be able to accommodate the N276 glycan by L-CDR1 point mutations based on similarity with bnAb lineage PCIN63 (*34*). HuGL17 and HuGL18 have an L-CDR3 sequence of C<u>QQYET</u>F, only 1 aa different than mature VRC01. HuGL18 was chosen for its high affinity (125 nM) and the use of VK3-20, a commonly used VK carrying a relatively short L-CDR1 of 7 aa. HC knock-in mice were generated using ES cell targeting (*12, 47, 48*). Knock-in allele usage for HuGL16, HuGL17 and HuGL18 was 89%, 79% and 93%, respectively, as determined in crosses with IgH^a/a^ wild type strains (**Fig S1A**, similar to the 85% usage in VRC01^gH^ mice (*12*). LC knock-ins were generated directly in zygotes using CRISPR/Cas9 technology using either a single-cut strategy for HuGL18 (**Fig 1B**) or a double-cut strategy for HuGL16 and HuGL17 (**Fig 1C**). As is commonly seen in such models, only a fraction (∼30%) of cells in LC-only mice retained expression of the knock-in allele (**Fig S1B**). When bred together, HuGL16 H/L, HuGL17 H/L and HuGL18 H/L mice (“HuGL16”, “HuGL17”, and “HuGL18” hereafter) generated 20%, 40% and 40% eOD-GT8-binding cells, respectively, as measured for HuGL17 and HuGL18 using eOD-GT8:streptavidin or, for HuGL16, by binding of eOD-GT8 60mer nanoparticles (**Fig 1F,G**). B cell numbers were somewhat lower than normal C57BL/6 B cell numbers (**Fig S1**), comparable to many BCR knock-in models. A significant proportion of eOD-GT8-binding cells in the bone marrow, spleen and lymph nodes of HL mice expressed only the knock-in chains, however some cells co-expressed an additional endogenous LC (**Fig 1D, Fig S1D,E**). Importantly, eOD-GT8-binding cells were present in similar percentages in spleen and lymph nodes, indicating they were likely functional, as lymph nodes are enriched in long-lived, non-anergic naive B cells (**Fig 1E-H, S1F-G**). B cells from HuGL17 and HuGL18 were able to make robust calcium flux responses to eOD-GT8 nanoparticles, whereas HuGL16 B cells failed to respond detectably under these conditions (**Fig S2**). Epitope-mutated negative control nanoparticles (eOD-GT8-KO 60mer) failed to elicit any response (**Fig S2**). These three HuGL mouse models thus generated B cells with properties predicted to allow response to eOD-GT8, however the in vitro Ca^++^ response of low affinity HuGL16 B cells was poor.

**Figure 1.**
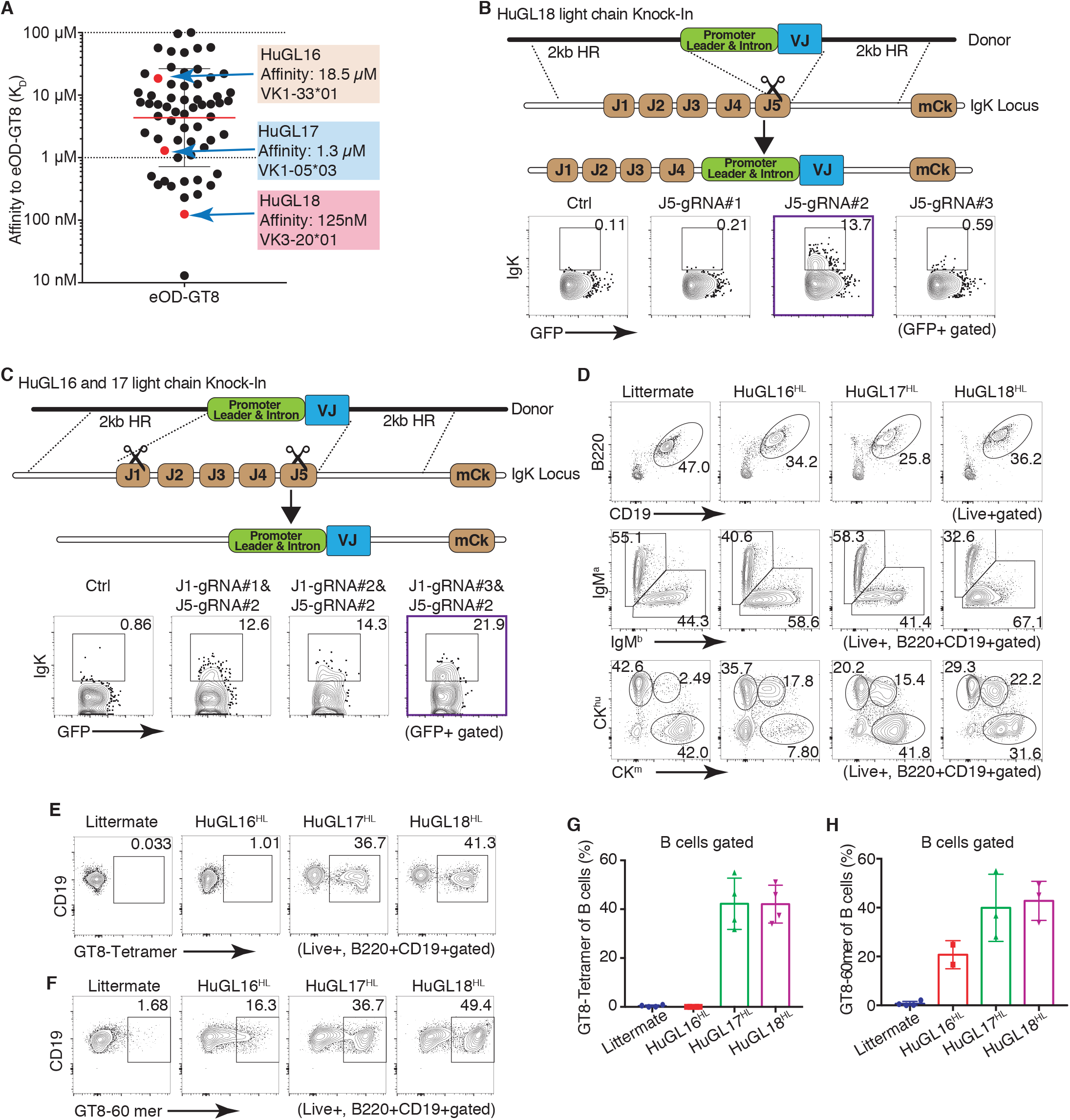
Naive human B cell BCR VRC01-class H/L knock-in mouse models HuGL16, HuGL17 and HuGL18. Figure shows the design and features of HuGL knock-in mice. (**A)** Affinities of VRC01-class BCRs cloned from HIV^−^ healthy human donors (*43*) highlighting the K_D_ values and LC usage of HuGL16, HuGL17 and HuGL18. (**B,C)** HC knock-ins were generated using a conventional targeting strategy as described (*12, 47, 48*). LC knock-in strategies are shown in the schematics, detailing the CRISPR-facilitated targeting strategy used for HuGL18 (**B**) or HuGL16 and HuGL17 (**C**), which involved one or two cuts in the Jκ locus, respectively. HR indicates arms of homology. Flow plots below each schematic show transient transfection analysis of gene targeting efficiencies in a *Rag1*^−/−^ pro-B cell-line using CRISPR ribonucleoproteins carrying the indicated guide RNAs and LC targeting construct (see Methods). (**D-H)** Characterization of lymphocytes in HuGL mice. (**D**) Alleles were marked by breeding BCR knock-ins of interest (C57BL/6 background: *Igh*^b/b^*C*κ^m/m^) to *Igh*^a/a^*C*κ^hu/hu^ mice, followed by flow cytometry analysis with antibodies that distinguish C-regions. (**E-H)** Evaluation of antigen-binding by B cells of the indicated strains using eOD-GT8 streptavidin tetramers (**E**) or eOD-GT8 60mer nanoparticles (**F**), with quantification shown in G,H. Data are representative of multiple litters per HuGL. (E-H) N = individual experiments, n = mice per group. N = 3. n = 1-3 mice per experiment. See also Supplementary Figure 1.

### HuGL18 VRC01-class B cells can be primed and recruited to GCs in vivo

We next tested whether these B cells carrying authentic naive human VRC01-class BCR specificities could respond functionally in vivo, to confirm that the germline-targeting immunogen “bait” used to isolate VRC01-class human naive B cells can not only bind intended BCRs *ex vivo* but can activate these cells in vivo. It was also of substantial interest to know whether VRC01- class HuGL B cells would be capable of participating in GCs. We first tested HuGL18 B cells. We transferred a high number of HuGL18 B cells (CD45.2^+^ allotype) into CD45.1^+^ congenic recipients, establishing a 1 in 1000 HuGL18 B cell precursor frequency, and immunized the recipients with eOD-GT8 60mer in alum. Env CD4bs-specific IgG responses were detectable in HuGL18 recipients by day 8 post-immunization (**Fig 2A**). CD4bs-specific IgG responses were not detected in mice immunized with eOD-GT8-KO 60mer (**Fig 2A**). After immunization with eOD- GT8 60mer, large numbers of HuGL18 B cells were detected in GCs by flow cytometry (CD38^−^ GL7^+^) (**Fig 2B-C**). As expected, these HuGL18 B cells were antigen-specific (**Fig 2D**). Immunofluorescence staining of spleen histology sections revealed numerous GCs that contained HuGL18 B cells (**Fig 2E**), in agreement with the flow cytometric data. Taken together, the data show that B cells expressing authentic VRC01-class human naive B cell BCRs are functional in vivo.

**Figure 2.**
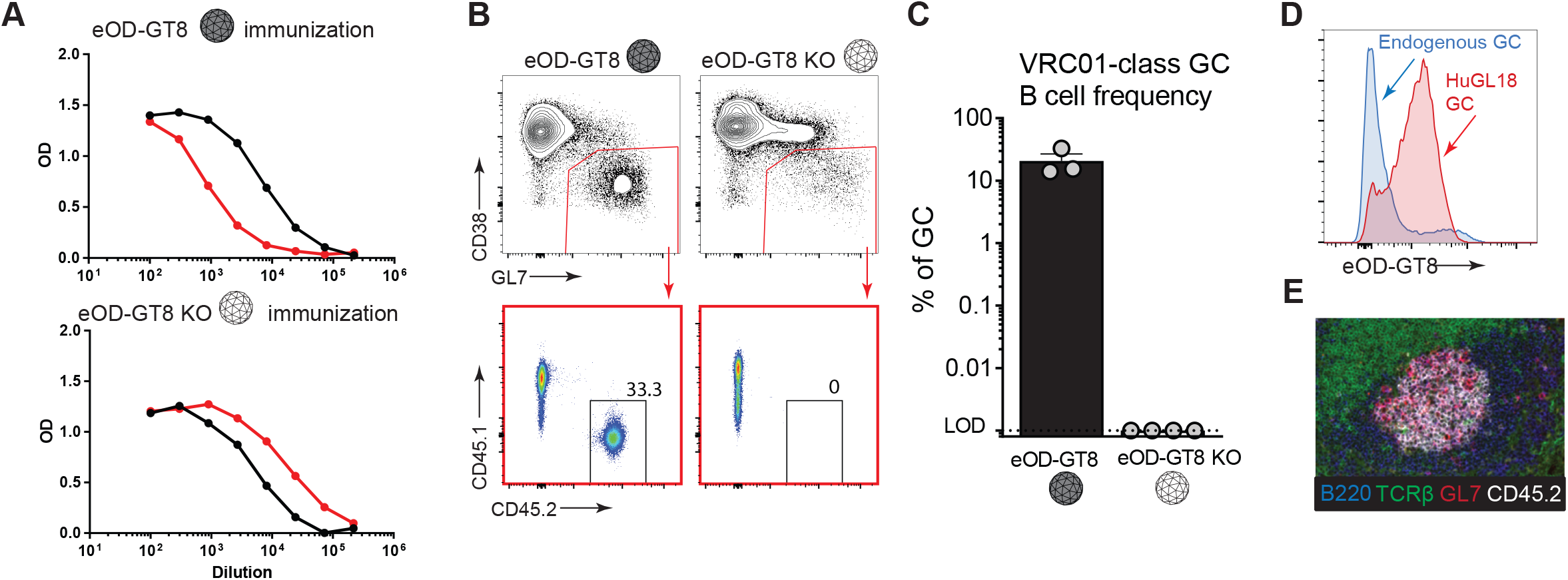
VRC01-class B cells from a BCR knock-in mouse with an authentic naive human VRC01-class BCR specificity (HuGL18) can be primed and recruited to GCs. **(A)** Day 8 ELISA binding curves from CD45.1^+^ mice that received HuGL18 B cells (1 in 10^3^ precursor frequency) and were immunized with eOD-GT8 60mer (top) or eOD-GT8-KO 60mer (bottom). eOD-GT8 IgG titers are shown in black, KO IgG titers are shown in red. **(B)** Flow cytometric analysis of total B_GC_ (gated as SSL/B220^+^/CD4^-^/CD38^-^/GL7^+^) and HuGL18 B cell occupancy of GCs (SSL/B220^+^/CD4^-^/CD38^-^/GL7^+^/CD45.1^-^/CD45.2^+^ cells) of mice immunized with eOD-GT8 60mer (top), or eOD-GT8-KO 60mer (bottom). Day 8 splenic B cells were analyzed. **(C)** Quantitation of VRC01-class B cells among B_GC_ cells. **(D)** Flow cytometric analysis of antigen-binding (eOD-GT8) by endogenous or HuGL18 B_GC_ cells. Representative of 5 mice tested. Mice immunized as in (B). **(E)** Histological analysis of GCs. Frozen splenic sections were stained with B220 (blue), TCRβ (green), GL7 (red), and CD45.2 (white) for identification of HuGL18 B cells within GCs. SSL=Singlet Scatter Live. N = 3 (B-C), N = 1 (A,D,E). n = 3-4 mice per experiment.

### HuGL18 VRC01-class B cells at rare physiological precursor frequencies can be primed by eOD-GT8 60mer and recruited to GCs

In vivo validations of germline-targeting approaches to vaccine design are key, particularly under physiological relevant conditions, given that bnAb precursor B cells are rare in humans (*28, 29, 43*). It is not uncommon for in vivo mouse models to utilize precursor frequencies of 1 in 10 B cells to 1 in 1000 B cells, which are far higher than the physiological range of a normal repertoire (*45, 49*). We therefore next assessed if HuGL18 VRC01-class B cells could be primed at physiologically relevant precursor frequencies found in the human repertoire. Human naive VRC01-class B cells with affinities better than a K_D_ of 3 μM (the top ∼33%) are found at a precursor frequency of approximate 1 in 10^6^ B cells (*43*). We hypothesized that precursor frequency would be an important determinant for recruitment of authentic HuGL18 B cells to early GCs. To test the hypothesis, we transferred a range of HuGL18 B cells into congenic CD45.1^+^ hosts and immunized. The spleens of mice receiving HuGL18 B cell transfers were assessed for grafting efficiency, and precursor frequencies of 1 in 10^3^ B cells to 1 in 10^6^ B cells were confirmed (data not shown). On day 8 post-immunization with eOD-GT8 60mer, all HuGL18 recipient animals developed similar total GC B cell (B_GC_) frequencies **(Fig 3A)**. However, the frequency of HuGL18 B cells among B_GC_ cells was sharply dependent on the original precursor frequency. When HuGL18 B cells started from high precursor frequencies (1 in 10^3^-10^4^), HuGL18 B cells occupied a high proportion of the B_GC_ compartment (∼10-40%, **Fig 3B-C**). Contrary to this, when precursor frequency was restricted to the physiological level found in humans (1 in 10^6^ B cells), HuGL18 B cell frequencies among B_GC_ post-immunization were 40-160-fold lower (∼0.25% of the GC compartment, **Fig 3B-C**). The HuGL18 cells localized to GCs histologically, consistent with the frequencies determined by flow cytometry **(Fig 3D)**. The HuGL18 B cell response was also confirmed to be CDbs-epitope-specific; immunization with mutant eOD-GT8-KO 60mer did not prime HuGL18 B cells (**Fig 3B-C**). We next sought out how HuGL18 GC B cells could compete within the GC compartment over time when starting from physiologically rare precursor frequencies of 1 in 10^6^ B cells. HuGL18 GC B cells showed outgrowth in most animals over time, achieving upwards of 5.7% of B_GC_ cells at day 20 post-immunization **(Fig 3E)**. In summary, B cells expressing authentic VRC01-class human BCRs could be primed to proliferate and be recruited to GCs by the eOD-GT8 60mer germline-targeting antigen, even when the HuGL B cells start from rare physiologically relevant precursor frequencies.

**Figure 3.**
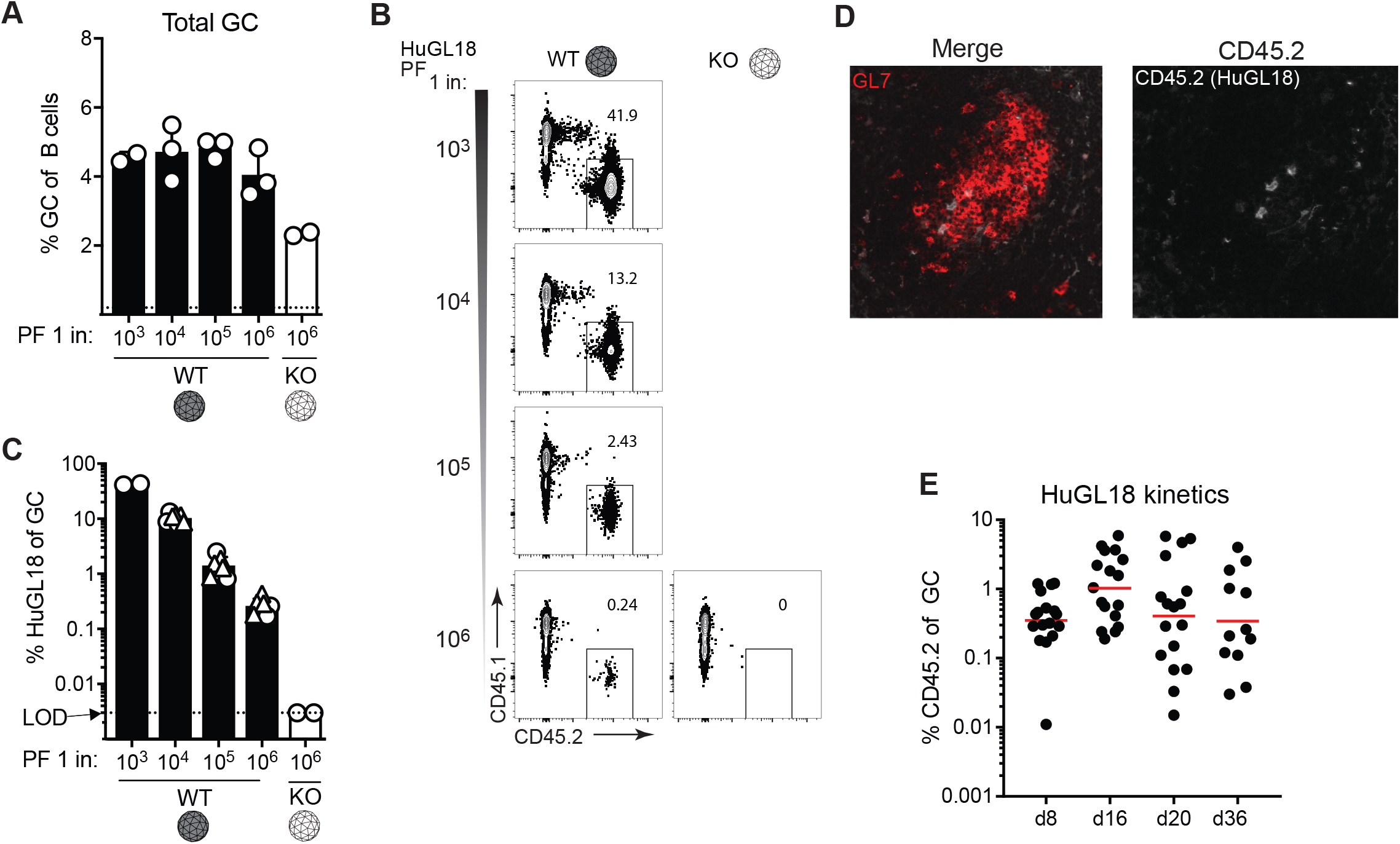
Authentic human VRC01-class naive B cell BCRs can be primed by eOD-GT8 60mer and recruited to GCs at rare physiological precursor frequencies. **(A)** Frequency of splenic B_GC_ cells in mice immunized with eOD-GT8 60mer, or the eOD-GT8- KO variant, when starting from the indicated HuGL18 B cell precursor frequency (PF). Day 8 splenocytes were analyzed and B_GC_ cells were gated as SSL/B220^+^/CD4^-^/CD38^-^/GL7^+^ and plotted as % of total B cells (B220^+^ cells). **(B)** Representative flow cytometric plots of HuGL18 B cells in GCs utilizing allotype mark. Cells are gated on GCs as indicated in A. **(C)** Quantitation of HuGL18 B cells in GCs as indicated in B. Experiments are pooled between two experiments (circles and triangles are separate experiments). 25 total mice examined. **(D)** Histological analysis of individual GCs for HuGL18 B cells in individual GCs as indicated in B. N = 3. n = 3-4 mice per experiment.

### HuGL18 B cells develop memory after eOD-GT8 60mer immunization

A bnAb-based HIV vaccine approach will involve multiple booster immunogens, designed to recruit VRC01-class memory B cells back to GCs and “shepherd” their affinity maturation pathway toward broad neutralization (*9, 14, 25*). We therefore evaluated VRC01-class memory B cell development after an eOD-GT8 60mer priming immunization, starting from physiologically rare naive VRC01-class B cell precursor frequencies. All immunized mice developed HuGL18 memory B cells by day 36 post-immunization **(Fig 4A)**. The frequency of memory B cells was variable between mice and achieved frequencies of 1 in 10^5^-10^6^ B cells **(Fig 4B)**. Nearly all of the HuGL18 memory B cells were class switched **(Fig 4C-E)**. Additionally, the majority of HuGL18 memory B cells expressed CD73, PD-L2, and CD80 **(Fig 4F-G)**, which are three memory B cell markers associated with GC-derived memory B cells (*50, 51*). Altogether, the data show that HuGL B cells can form GC-derived memory B cells, even when starting from physiologically rare precursor frequencies.

**Figure 4.**
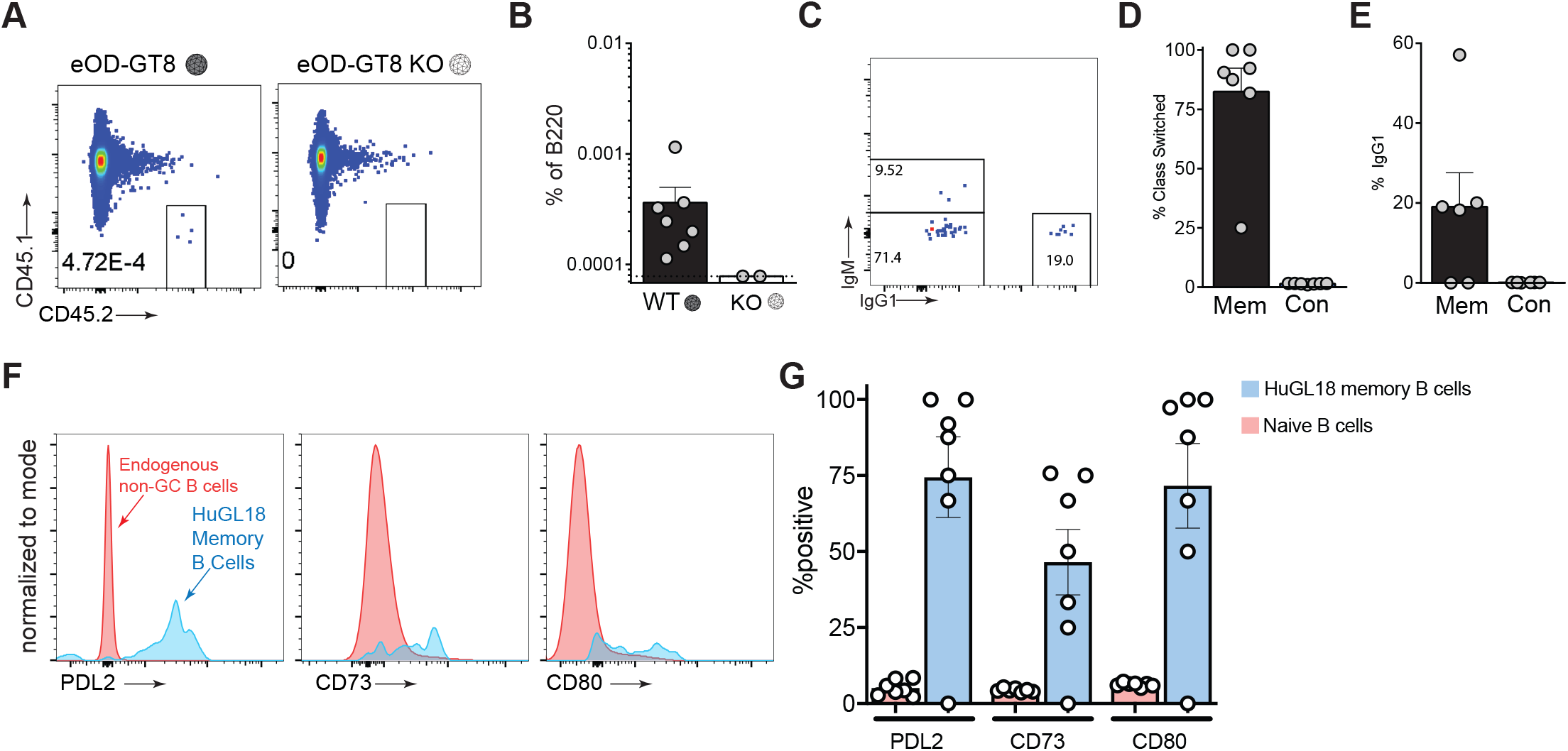
HuGL18 B cells develop memory after eOD-GT8 60mer immunization. **(A)** Mice were assessed by flow cytometry for HuGL18 memory B cell formation in the spleen on day 36 post-immunization with eOD-GT8 60mer. eOD-GT8-KO 60mer immunization was done as a negative control. Precursor frequency of HuGL18 naive B cells was 1 in 10^6^ before immunization. Memory B cells were gated as SSL/B220^+^/CD4^-^/IgD^-^/CD38^+^/GL7^-^/CD45.2^+^/CD45.1^-^. **(B)** Quantitation of memory B cells in A, as % of B220^+^ B cells. **(C)** Flow cytometric analysis of class switched memory B cells gated in A. **(D-E)** Quantitation of (D) HuGL18 class switched memory B cells and (e) HuGL18 IgG1^+^ memory B cells on day 36 post immunization. **(F-G)** Memory B cells phenotypes. **(F)** Representative flow cytometry and **(G)** quantitation of indicated memory B cell markers on memory HuGL18 B cells on day 36. Red histograms are total non-B_GC_ cells (SSL/B220^+^/CD4^-^/CD38^+^/GL7^-^). N = 3. n = 4-7 mice per experiment.

### Vk3-20^+^ VRC01-class B cells develop SHM and affinity maturation after a priming immunization

HuGL18 VH1-2*02 HCs accumulated substantial aa SHMs within 16 days post-immunization, with a maximum of 11 aa mutations, a median of 3 aa mutations, and with >95% of HuGL18 B_GC_ clones containing at least 1 aa mutation (**Fig 5A)**. Substantially more SHM accumulated in HuGL18 B_GC_ cells by d36 (**Fig 5A-D**). SHMs in H-CDR2 were present **(Fig 5B-C, Fig S3A)**, consistent with the H-CDR2 dominant interaction with eOD-GT8 (*28*); and substantial mutations were also present in H-CDR1 and H-CDR3. SHM accrual in the H-CDR3 of HuGL18 was particularly prominent when compared to VRC01^gHL^ **(Fig 5B-C)**. This illustrates a value of assessing B cells with authentic naive H-CDR3s. Nearly all VRC01-class bnAbs contain a W at the W100b (Kabat numbering) equivalent position of VRC01 in the H-CDR3, though one bnAb possesses a Y. Naive human VRC01-class B cells generally possess a Y or W at this position (*43*). Mutations were observed in HuGL18 B_GC_ cells at Y107 (100b), including Y→P or Y→W (**Fig S3A**). VH1-2 positions known to be important for VRC01-class broad neutralization capacity were mutating, including M34L/I in H-CDR1, as well as K63Q/R and Y95F **(Fig 5B-C, Fig S3A)** (*9, 34*). We next determined how many VRC01-class mutations existed in each HC sequence (excluding the H-CDR3, which cannot be computationally assessed) and compared that to the total overall mutations to ascertain whether HuGL18 B cells were capable of maturing along a desired affinity maturation pathway in response to eOD-GT8 60mer. Encouragingly, we observed substantial directionality in the mutational pathway of HuGL18 B cells in GCs over time, in response to a single immunization **(Fig 5D)**. Some HuGL18 clones at d16 achieved a perfect or near VRC01-class mutational trajectory **(Fig 5D)**, while >77% of clones contained one or more VRC01-class mutation by d36.

**Figure 5.**
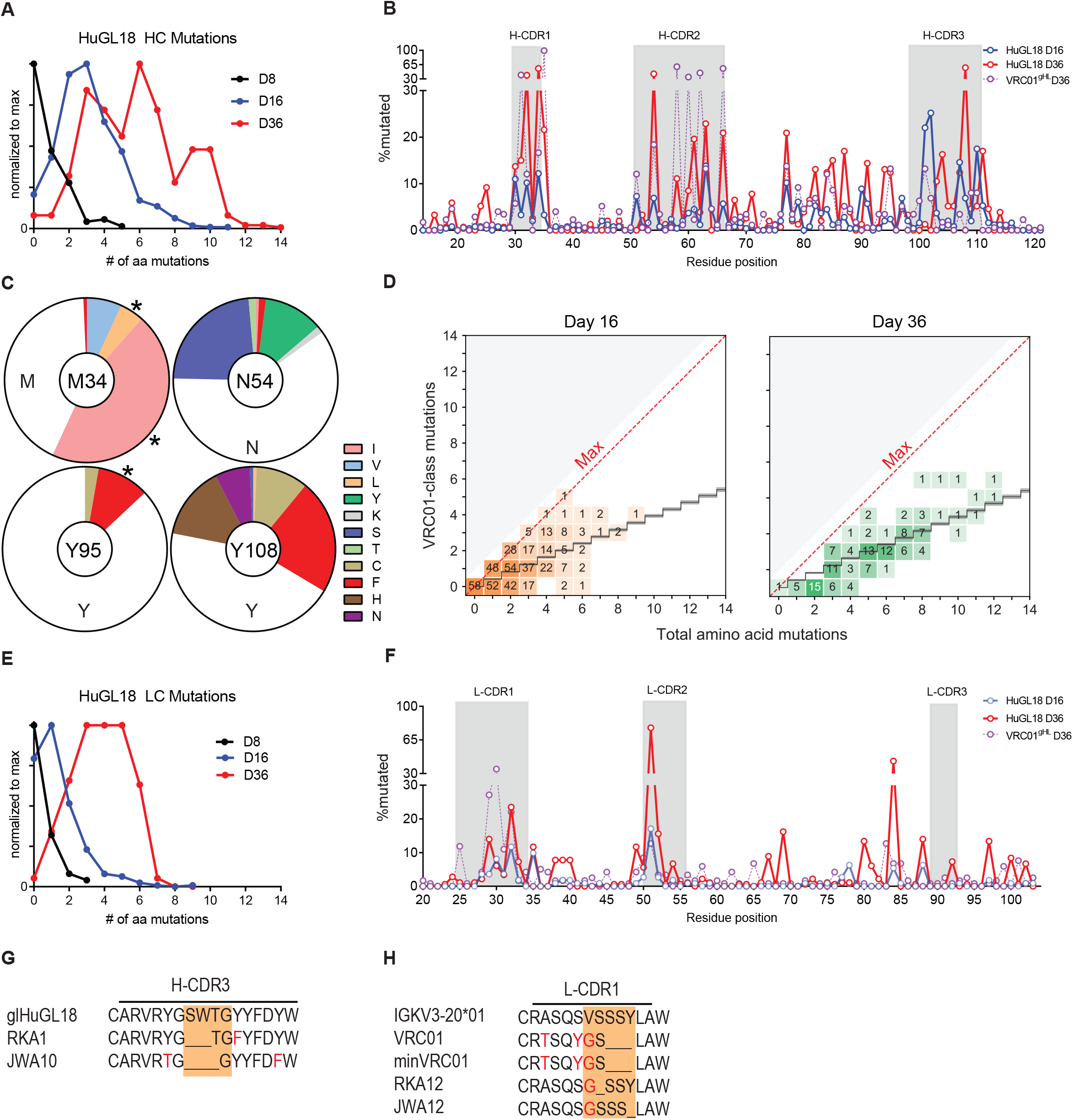
Vk3-20^+^ VRC01-class B cells develop SHM and affinity maturation after a single priming immunization. Mice were generated with a HuGL18 B cell precursor frequency of 1 in 10^6^ B cells, then immunized with eOD-GT8 60mer. Splenic IgG1^+^ d16 HuGL18 B_GC_ cells were sorted from eOD- GT8 60mer immunized mice on days 8-36 post immunization and HCs **(A-D, G)** and LCs (**E-F, H**) were assessed for aa SHMs. Cells were gated as SSL/B220^+^/CD4^-^/CD38^-^/GL7^+^/CD45.2^+^/CD45.1^-^/IgG1^+^/IgD^-^/CD138^-^. **(A)** Distribution of HC mutations over time. S = individual sequences. s = 134 for d8, s = 455 for d16, s = 148 for d36. **(B)** HuGL18 HC aa SHM distribution. For comparison, d36 VRC01^gHL^ SHM data from animals with 1 in 10^6^ precursor frequencies immunized with eOD-GT5 60mer (from (*27*)) are overlaid. **(C)** Specific HuGL18 HC aa mutations at d36. Asterisks (*) mark VRC01-class mutations. **(D)** VRC01-type mutations detected in HuGL18 HCs. A VRC01-class mutation (y-axis) is any mutation observed in a VRC01- class bnAb. The black staircase depicted is a computational estimate of antigen-agnostic mutation accumulation in VH1-2 B cells (*46*). **(E)** Total LC aa mutations. s = 72 for d8, s = 343 for d16, s = 179 for d36. **(F)** Per residue HuGL18 LC aa SHM distribution. D36 VRC01^gHL^ data are shown for comparison, as in B. (**G)** Alignment of two representative HuGL18 clones recovered on d36 with deletions in H-CDR3, out of 7 total clones recovered with deletions in H-CDR3. **(H)** Alignment of two representative HuGL18 clones recovered on d16 with deletions in L-CDR1, out of 13 total clones recovered with deletions in L-CDR1. Total sequences are from all experiments pooled. N = 2 for d16/d36. N=1 for d8. n=3-7 mice per experiment. See also Supplementary Figure 3.

The majority of HuGL18 B_GC_ cells at day 16 post-immunization also possessed LC mutations, with up to 9 aa SHMs detected **(Fig 5E).** Minimal mutation was seen in L-CDR3 (**Fig 5F)**, which only differed from mature VRC01 by one aa, thus preserving the critical VRC01-class features of the L-CDR3. Most mutations were in L-CDR1 **(Fig 5F)**, consistent with the presence of substantial affinity maturation in L-CDR1 of VRC01-class bnAbs.

Deletion events, which are often considered rare, were also observed in HuGL18 B_GC_ cells, in HC or LC. Seven HuGL18 B_GC_ clones, isolated from two separate mice, showed deletions of 2- 3 aa in H-CDR3 on d36 post-immunization (**Fig 5G**), which may be involved in the maturation pathway of VRC01 (*19*). Strikingly, 13 clones, isolated from two independent mice, contained single aa deletions in L-CDR1 **(Fig 5H).** Interestingly, 12 of these deletions were of S30 and one deletion was found in Y33, which aligns with the deletion found in the mature VRC01 bnAb and a minimally mutated VRC01 bnAb construct (*9*). Interestingly, all of the clones that contained deletions had an accompanying V29G mutation. V29G is present in the mature VRC01 and appears to be important based on the minimally mutated VRC01 bnAb construct (*9, 34*). Both of these observations are of considerable interest, as L-CDR1 deletions are critical for most VRC01-class Abs to develop breadth, to circumvent clashing with the Env N276 glycan. Deletions in L- CDR1 have been considered to be a major hurdle for VRC01-class bnAb vaccine development (*5, 19*). Here, we show that L-CDR1 deletions can occur in a HuGL B cell in as few as 16 days following eOD-GT8 60mer immunization.

In summary, authentic VRC01-class naive B cells primed at physiologically rare precursor frequencies were able to undergo substantial SHM in both the HC and LC and accrue specific VRC01-class mutations, including deletions in L-CDR1, in response to a single priming immunization. Thus, germline-targeting immunization strategies can both prime and affinity mature rare cells possessing epitope-specific naive human B cell BCRs, with a single immunization.

### A medium affinity, Vk1-5^+^ VRC01-class BCR knock-in mouse model

Immune responses by Vk1-5^+^ VRC01-class naive B cells are of specific interest because the first Vk1-5^+^ VRC01-class bnAb was recently identified, and that bnAb (PCIN63) has very appealing characteristics of low SHM% and the absence of any indels (*34*). Using the adoptive transfer approach, we assessed the immune response of HuGL17 B cells (Vk1-5^+^) (**Fig 6**). At a precursor frequency of 1 in 1000, a strong GC response of the donor cells was elicited by eOD-GT8 60mer but not the eOD-GT8-KO 60mer (**Fig 6A**). Responding donor B_GC_ cells underwent a robust IgG1 class switch comparable to that of the host response (**Fig 6B**). We then assessed the response of HuGL17 cells seeded at a precursor frequency of 1 in 10^6^ (**Fig 6C-I**). The total (host + donor) GC response peaked at ∼d8 (**Fig 6C**). The HuGL17 cells proved able to participate in the GC response (**Fig 6D**), and HuGL17 B_GC_ cells comprised an increasing fraction of the GC response through d21 (**Fig 6E**). The HuGL17 VRC01-class B cell response plateaued at approximately 0.1% of the total B_GC_ and 0.005% of total B cells, representing a ∼50-fold expansion from input on average (**Fig 6F**). The variable nature of the outgrowth of HuGL17 B cells was observed across experiments. At this low precursor frequency, we were unable to consistently enumerate HuGL17 memory B cells, but HuGL17 memory B cells were consistently observed in experiments starting from a precursor frequency of 1 in 10^5^ (**Fig S4**). These data indicate that, when primed by eOD-GT8 60mer, Vk1-5^+^ VRC01-class naive B cells were able to participate and persist in the highly competitive GC response under physiological precursor frequency conditions.

**Figure 6.**
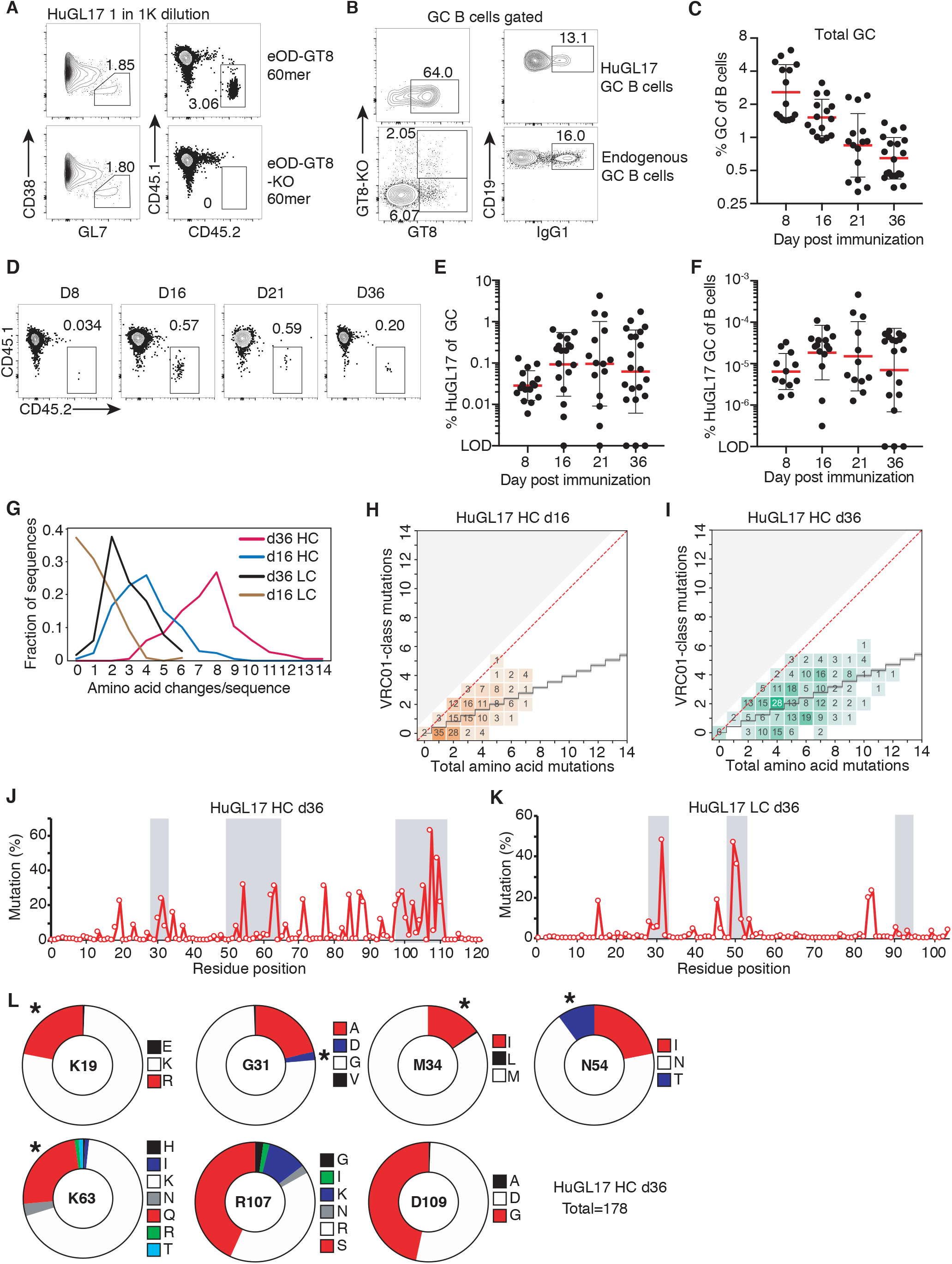
Characterization of HuGL17, a medium affinity, VK1-5^+^ VRC01-class BCR model. Analysis of total and HuGL17 splenic B_GC_ cells. (**A,B**) Analysis of HuGL17 responsiveness to eOD-GT8 in vivo at a HuGL17 B cell precursor frequency of 10^−3^ (“1K” = 1,000). (**A**) Total (left panels) and donor-derived fraction (right panels, boxes) of B_GC_ cells after immunization with eOD- GT8 or eOD-GT8-KO. (**B**) Representative flow plots of antigen-specific and IgG1 class-switched HuGL17 and endogenous (CD45.1^+^) B_GC_ cells (TCRβ^−^CD19^+^CD38^−^GL7^+^). (**C-L**) Longitudinal analysis of mice immunized with eOD-GT8 60mer when HuGL17 precursor frequencies were 1 in 1 million B cells. (**C**) Longitudinal analysis of total frequency of B_GC_ cells. (**D**) Representative flow plots enumerating HuGL17 B_GC_ cells. (**E**) Longitudinal analysis of GC occupation by HuGL17 B cells. Each data point indicates the value measured in one recipient spleen. (**F**) HuGL17 B_GC_ cell % among all B cells. **(G)** Analysis of the number of aa replacements per sequence in the HC and LC, respectively, at d16 (n=197, 172) and d36 (n=178, 226). **(H,I)** Analysis of HC VRC01-class mutations on d16 and d36. (**J,K**) Distribution of replacement mutations as a function of aa position. **(L)** Quantitation of HC replacement mutations d36 (n = 178). Asterisks (*) mark VRC01-class mutations. N = 3. n = 2-7 mice per experiment.

Antibody gene sequencing analysis of HuGL17 B_GC_ cells at d16 and d36 revealed extensive SHM, particularly in the HC (**Fig 6G-L**. **Fig S5A)**. Many HC mutations within VH1-2 were identical to mutations seen in VRC01-class bnAbs (**Fig 6H,I,L**. **Fig S5C)**. The L-CDR3 was barely mutated (**Fig 6K**. **Fig S5B)**, preserving the critical residues found in VRC01-class bnAbs. In contrast, the HuGL17 H-CDR3 was heavily mutated (**Fig 6J**. **Fig S5A**), especially at R107, an arginine lying at the center of the H-CDR3 loop that may be subject to negative selection. In the Vk1-5^+^ bnAb PCIN63, four glycine replacements are present in the L-CDR1 instead of the small L-CDR1 deletion observed in the bnAb VRC01. Although eOD-GT8 is not designed to specifically select for VK1-5 L-CDR1 features, six HuGL17 B_GC_ cells were found to have introduced glycine replacement mutations in L-CDR1, which is predicted to facilitate evolution to neutralization breadth. One HuGL17 B_GC_ clone contained a doubly glycine replacement in L- CDR1 positions identical to that of the bnAb PCIN63. We conclude that, at these physiological precursor frequencies, HuGL17 B cells participate in the GC response and undergo affinity maturation, indicating that the Vk1-5^+^ VRC01-class naive B cells identified in humans are likely targets, and interesting targets, for priming by eOD-GT8 60mer in humans.

### Precursor frequency and affinity are interdependent in determining competitive success for B cells possessing authentic human naive VCR01-class BCRs

There is evidence that both B cell precursor frequency and BCR affinity influence B cell competition following immunization (*24, 27, 45, 52, 53*). To quantitatively assess this prediction with B cells expressing authentic naive human BCR sequences, we transferred 10-fold serial dilutions of HuGL B cells into congenic recipients and immunized with eOD-GT8 60mer. These HuGL B cells not only utilize three LC V genes found in VRC01-class bnAbs (VK3-20, VK1-5, and VK1-33), but HuGL18, HuGL17, and HuGL16 also span a physiological range of affinities (0.126 μM→18.5 μM) found in humans. At day 8 post-immunization, a clear hierarchy in GC occupancy was observed based both on the starting precursor frequencies of the naive B cells and the BCR affinity **(Fig 7A-B)**. Combining the available data, the competitive fitness of high (HuGL18) and medium (HuGL17) affinity cells over time can be compared. Using experiments with HuGL18 or HuGL17 seeded in recipient mice at the 1 in 10^6^ physiological precursor frequency (**Fig 3E**, **Fig 6E**), the magnitude of the VRC01-class B_GC_ cell responses correlated with starting BCR affinity (**Fig 7C**). While HuGL18 B cells were present in GCs in higher numbers than HuGL17 as early as d8 (P < 0.001, **Fig 7C**), on average the HuGL17 B cells did shrink the gap with the HuGL18 B_GC_ response (d16, 10-fold difference; d21, 2-fold difference; d36, 5-fold difference. **Fig 7C**). HuGL18 cells exhibited rapid population of GCs by d8, calculated to be a ∼100-fold outgrowth of cells from the d0 HuGL18 precursor frequency. HuGL17 B cells showed greater outgrowth from d8-d16 within GCs (**Fig 7C**). This highlights that the affinity of precursors as well as Vκ gene usage may affect the numbers of initial GCs seeded and/or the kinetics within GCs. Taken together, these data highlight the importance of precursor frequency and antigen affinity in vaccine design, and this pre-clinical model predicts, with certain caveats, that the ongoing clinical trial of the germline-targeting HIV vaccine antigen eOD-GT8 60mer will prime desired human VRC01-class B cells and recruit those cells to GCs in a manner dependent both on B cell precursor frequency and antigen affinity.

**Figure 7.**
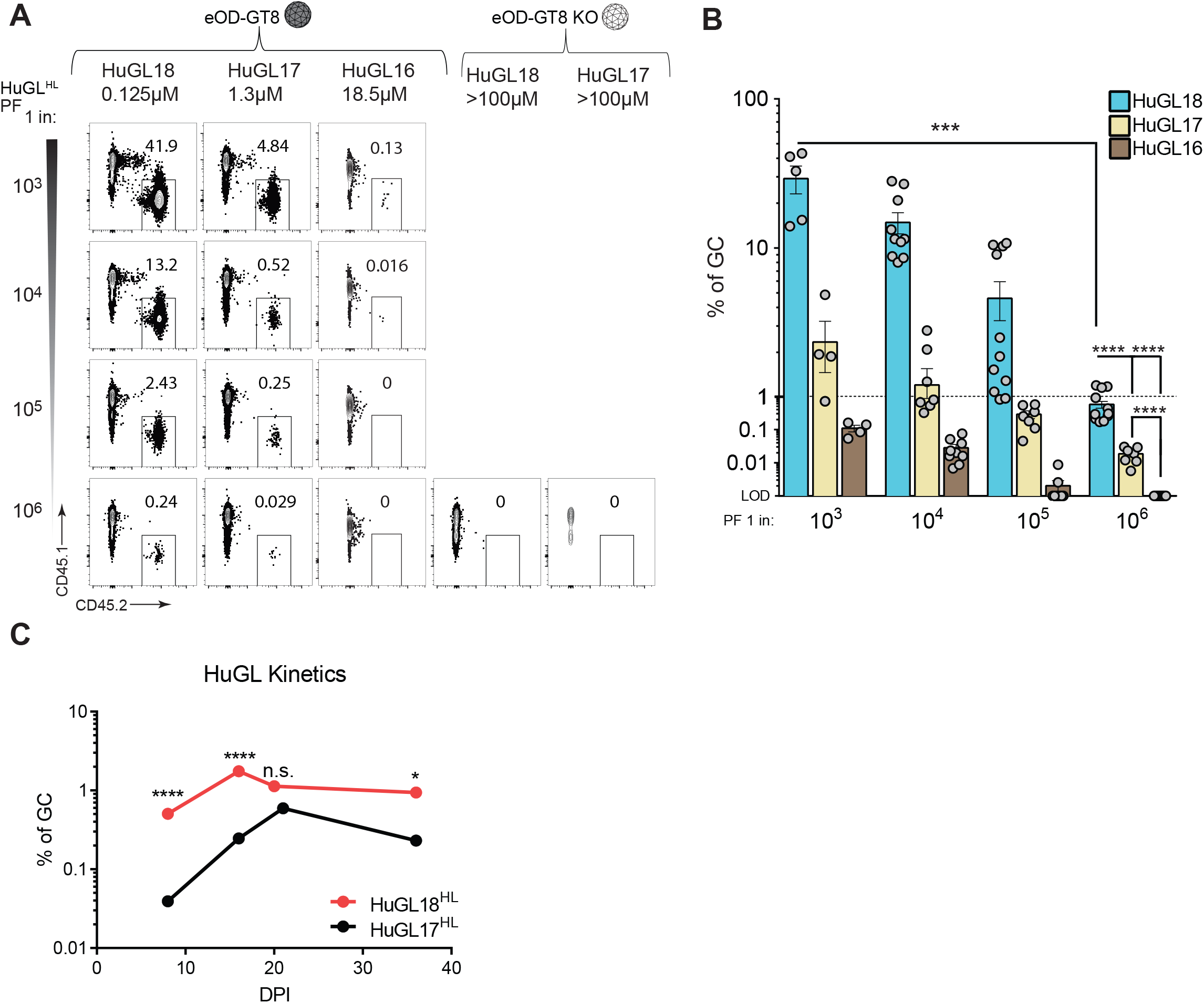
Precursor frequency and affinity are interdependent in determining competitive success for B cells possessing authentic human naive VCR01-class BCRs. **(A)** Frequency of HuGL B cells in GCs d8 post-immunization following immunization with eOD- GT8 60mer or eOD-GT8KO 60mer. B_GC_ cells gated as SSL/B220^+^/CD4^-^/CD38^-^/GL7^+^/ CD45.1^-^/CD45.2^+^. For comparison purposes between mouse strains, HuGL18 plots (Fig 3) are reshown here. **(B)** Quantitation of HuGL B cells in GCs as shown in A. All available data are shown. Data were pooled from multiple independent experiments. (**C)** Statistical comparisons of HuGL GC B cells over time. Kinetics data shown in 7C are the average of all data shown in Fig 3E and Fig 6E. Mice were immunized with eOD-GT8 containing HuGL17 or HuGL18 B cells at a precursor frequency of 1 in 10^6^ B cells. ** p<0.001, *** p<0.0001. N=3, n=3-4 mice per experiment for A-B. N=4-5 for HuGL17 and HuGL18 each, n=3-7 mice per time point for C.

## DISCUSSION

Here, for the first time, BCRs isolated from authentic antigen-specific naive human B cells were tested in mouse models for their ability to respond to antigen, somatically hypermutate, and compete in germinal centers. VRC01-class naive B cells from all three BCR H/L knock-in mouse models were able to be primed in vivo and participate in GCs. High affinity and medium affinity VRC01-class B cells (HuGL18 and HuGL17) were able to successfully compete in GCs and affinity mature—including acquisition of bnAb-type somatic mutations—even when the HuGL B cells started from rare physiological precursor frequencies.

Germline-targeting vaccine design is based on the concept of designing vaccine antigens to bind precursors of highly potent neutralizing antibodies isolated from infected humans. Cycles of antigen design and immunization strategy iteration are almost certainly required for germline-targeting immunogen designs to be applicable for human vaccine needs. One strategy is to design germline-targeting antigens based on iGL sequences from bnAbs and then use the germline-targeting immunogen to screen the human B cell repertoire for the capacity of the putative germline-targeting immunogen to identify (bind) bnAb precursors in normal healthy donors, selecting top candidate immunogens to move forward into clinical trials (*6, 7*). A significant gap in this strategy has been the absence of an in vivo model to demonstrate the in vivo specificity and responsiveness of the human antigen-specific naive B cell BCR sequences prior to a full clinical trial. One solution to this problem, employed here, is to express antigen-specific human naive B cell BCRs in mice and thus provide a direct preclinical platform for evaluating relevant antigen design and immunization strategy iterations. The approach of identifying antigen-specific human naive B cells and developing HuGL mouse models of preclinical vaccine testing is also applicable more broadly to other, non-germline-targeting, reverse vaccinology antigen design strategies.

We found precursor frequency and antigen affinity to be critical in dictating B cell outcomes following vaccination. We observed that these factors were interdependent in determining recruitment to GCs, competition within GCs, and exit to the memory B cell pool. These HuGL results were consistent with our previous study (*27*). While HuGL B cell competitive success in GCs past day 8 was more stochastic between individual mice than seen for VRC01^gHL^ iGL B cells, BCR affinity was a determining factor in competitive success of HuGL B cells over time. The inter-animal variance of HuGL GC B cell responses may be a strength of this model by being reflective of variance that may be seen in humans. This stochasticity reinforces the importance of interclonal competition and immunodominance as hurdles in vaccine development to complex antigens (*54–56*). As the precursor frequency of virtually all bnAb B cells is predicted to be low (*9*), the present study indicates that a pre-immune BCR K_D_ of >10 μM would be insufficient for efficient recruitment into the nanoparticle response, and that vaccine immunogens should be designed with considerably tighter binding to BCRs of authentic naive human B cell targets (0.1-1 μM or better).

A major goal of mouse immunology is to model human immune responses; however, mice and humans have substantially different immunoglobulin gene repertoires, limiting the ability to test reverse vaccinology design concepts outside of humans. While multiple mouse lines have been designed to express diverse human immunoglobulin genes, a limitation of those models is that they fail to properly represent the precursor frequencies found in humans. Precursor frequency can be a major factor in B cell responses, and immunoglobulin-locus transgenic animals generally have greatly underrepresented (*23*) or overrepresented B cells of interest (*11–13, 22, 25, 26, 46, 48, 57*). In contrast, transfer models such as those used here can titrate the cell numbers to match the physiological precursor frequencies found in humans (*27*). In addition to precursor frequency issues, BCR knock-in models can depend on iGL sequences of bnAbs (*12–14, 25, 26, 58*). Such models have key caveats for interpreting how a candidate antigen may perform in humans. First, the H-CDR3 and L-CDR3 generally remain unchanged from the somatically hypermutated bnAb (because the original CDR3s cannot be accurately predicted due to stochastically determined sequence variations generated in each naive B cell during development). This is generally expected to result in significant unintended advantage to those iGL BCR B cells compared to authentic naive B cell BCRs. Second, bnAb iGL sequences are not known to be reliable proxies for true naive B cell BCR sequences represented in most humans. Lastly, the bnAb iGL sequence in the mouse model is frequently an exact match to a bnAb iGL sequence used in the immunogen design. Here, we provide a more rigorous test. eOD-GT8 was designed based on a broad panel of bnAb iGLs (*12*). eOD-GT8 was then tested for its ability to ‘fish out’ epitope-specific human naive B cells (*28, 43*). To our knowledge, eOD-GT8 is only one of two engineered vaccine antigens to successfully pass the test of binding epitope-specific human naive B cells directly from multiple human blood samples (*28, 29, 43*). (The predecessor of eOD-GT8, eOD-GT6, was designed based on a more narrow panel of bnAb iGLs and did not successfully pass the test of binding VRC01- class precursor naive B cells in human blood samples (*28*), highlighting the magnitude of the protein design challenge for binding diverse authentic human B cell BCRs.) By isolating dozens of epitope-specific naive human B cells with eOD-GT8 (*43*), we were able to categorize and stratify the BCRs.

The use of CRISPR technology greatly accelerates the speed of BCR knock-in mouse generation at the HC locus (*59, 60*). In this work, we used CRISPR-facilitated targeting in zygotes to introduce functional knock-ins into the κ-locus for the first time, developing models for three distinct LCs along with their corresponding HC models.

Here, we selected three representative VRC01-class naive B cell BCRs for construction of BCR knock-in mice, in an effort to make a multi-pronged set of mouse models for predicting potential outcomes of germline-targeting vaccine immunization in humans. Based on the experiments reported herein, these models indicate that a human clinical trial with eOD-GT8 60mer should be successful, based on the parameter of activating and expanding VRC01-class naive B cells. Both high and intermediate affinity VRC01-class naive B cell BCRs successfully expanded upon eOD-GT8 immunization in HuGL transfer models, when HuGL B cells were present at a precursor frequency of 1 in 1 million. The available data indicate that high and intermediate affinity BCRs (< 3 µM) are present in the human repertoire at a frequency of ∼1 in 0.9 million. Low affinity VRC01-class BCR naive B cells are several fold more abundant, but, at least for HuGL16, that does not suffice to overcome the competitive disadvantage of the weaker affinity cells.

Interesting VRC01-class B cell SHMs were observed in this study using HuGL B cells. VRC01-class mutations were observed in HCs and LCs of HuGL18 and HuGL17 B cells after only a single immunization with eOD-GT8 60mer. The H-CDR3s in both HuGL18 and HuGL17 B cells underwent substantial SHM. While most VRC01-class bnAbs have limited engagement of HIV Env via H-CDR3, recent analysis suggests that development of the VRC01 bnAb and the related VRC08 bnAb involved extensive mutation of H-CDR3 that affected the development of breadth. Deletions are generally considered very rare events, and deletions in L-CDR1 are thought of as a major hurdle to overcome in the development of VRC01-class bnAbs. Intriguingly, we found that deletions in L-CDR1 of HuGL18 B cells could be found in B_GC_ cells in as little as 16 days after eOD-GT8 60mer immunization. Taken together, these data highlight the importance of studying B cell responses by B cells expressing human BCRs from authentic naive B cells.

It remains unclear how to induce a full VRC01-class bnAb maturation path by immunization, and it remains unclear what fraction of VRC01-class naive B cells could successfully navigate such a path under optimal conditions. VRC01-class B cells refers to all B cells with the core HC and LC sequence characteristics of VRC01-class bnAbs: VH1-2*02 [or *03, or *04] paired with a LC possessing a 5 aa CDR3. A subset of naive B cells with those characteristics bind to eOD-GT8, generally reflecting clearer relatedness to VRC01-class bnAbs (i.e., L-CDR3 with a QQYxx motif and a preference for a short L-CDR1) (*9, 43*). Development of a VRC01-class B cell into a bnAb over time is not a given, as even in HIV^+^ donors with identified VRC01-class B cell responses, lineages or branches of lineages can fail to develop breadth (*19, 32, 61*). Whether some VRC01-class naive B cells are more likely to develop breadth remains an important open question for future investigation. For example, data shown here indicate that both VK3-20^+^ and VK1-5^+^ VRC01-class B cells have the ability to rapidly acquire mutations in L- CDR1 that are frequently considered important for accommodating Env N276. In the case of VK3- 20^+^, L-CDR1 deletions were detected among B_GC_ cells. In the case of VK1-5^+^, a pair of glycine mutations were found in L-CDR1. Previously it was thought that such mutations may be extremely rare. Lastly, the recently identified VK1-5^+^ VRC01-class bnAb PCIN63 lineage acquired breadth with as little as 19% aa mutations in the HC (*34*). It was encouraging to observe up to 12.7% aa mutations in both HuGL18 and HuGL17 B_GC_ cells after only a single immunization with eOD- GT8 60mer (**Fig 5**, **Fig 6**).

The HuGL models have limitations. The best assessment of those limitations will come with the completion of the full eOD-GT8 60mer clinical trial (the trial is currently ongoing). Human adaptive immunity cell biology is not identical to that of mice, and the B cell repertoires of the species are different, resulting in differences in the competitive environment that cannot be calculated and may or may not be sufficiently different to impact post-immunization outcomes. Overall, human versus mouse differences could theoretically cause a range of differences in the outcomes between mouse and human immunization with a germline-targeting antigen. Given the theoretical nature of those differences, we feel that the most productive current use of HuGL mouse models is to make specific predictions, allowing for clear assessment of whether specific predictions were useful, accurate, or inaccurate when the full clinical trial results become public. At least six different mouse models of VRC01-class B cell responses to eOD-GT have been tested, including the HuGL model reported here (*11, 22, 23, 27, 46*). It will be of great value to compare each of the mouse models to the human clinical trial results. Experimental advantages of other models include the use of truly polyclonal antigen-specific repertoires (*22, 23*), or the presence of high precursor frequencies or affinities making positive results in the model more likely (*11–13, 46*), or the use of the prototypic VRC01 iGL (*11, 12, 27, 46*).

The HuGL approach reported here is the only mouse model utilizing exact naive BCR sequences known to exist in the naive B cell repertoires of HIV-negative humans with truly authentic H-CDR3s. It is also the only model to match the known precursor frequency and affinity range in humans. The HuGL strategy has the added advantage of straightforward cell transfers to modulate the precursor frequencies, unlike models using genetically modified mice in a mixed background (*22, 23*). It is conceivable these HuGL may underestimate how well VRC01-class B cells can respond in humans due to challenges of introducing human BCR sequences into mice, as we observed that a fraction of transgenic B cells in all three mice co-expressed an endogenous mouse LCs and had somewhat reduced overall BCR levels. Lastly, the experiments demonstrate the power of new CRISPR-based approaches to make HuGL-type mouse models, because of the speed of new mouse generation.

Overall, these data indicate that VRC01-class B cells with VK3-20 or VK1-5 are likely to respond to eOD-GT8 immunizations of humans, and the data imply that VRC01-class B cells of any recognized subtype (VK3-20, VK1-5, VK1-33, VK3-15, VK1-4) are likely to respond in humans in a manner that is interdependent of precursor frequency and affinity. Furthermore, the HuGL mouse model data predict that the expansion of high and intermediate VCR01-class B cells should be accompanied by substantial SHM and possibly rare events such as deletions, even after a single immunization. These data complete a key loop of germline-targeting preclinical development strategy, and this strategy is generalizable to other germline-targeting and reverse vaccinology strategies for HIV or other pathogens.

## ACKNOWLEDGEMENTS

### Funding

This work was supported in part by the NIH NIAID under awards Al100663 (Scripps Center for HIV/AIDS Vaccine Immunology and Immunogen Discovery) and UM1 AI144462 (Scripps Consortium for HIV/AIDS Vaccine Development) (to S.C, W.R.S.); NIH K99 AI145762 (R.K.A.) by the Ragon Institute of MGH, MIT, and Harvard (W.R.S.); and by the International AIDS Vaccine Initiative (IAVI) Neutralizing Antibody Consortium (NAC) and Center (W.R.S.); and through the Collaboration for AIDS Vaccine Discovery funding for the IAVI NAC Center (W.R.S.).

### Author Contributions

CHD and SC conceived of the study. CHD, RKA, DH, WRS, DN, and SC designed the study. RKA and DH performed immunization studies. DH, GM, SK constructed LC mice. DN, SAV, GM, SK, TCT constructed HC mice. WRS designed immunogens. PDS managed mice. NP, RT, and BG provided immunogens and sorting reagents. SK provided bioinformatic analysis and project management service. RKA, DN, and SC wrote the manuscript. Co-authors edited the manuscript. W.R.S. is involved as a PI of a human clinical trial involving eOD-GT8 60mer (G001). As part of that involvement, W.R.S. is privy to certain clinical trial data, and he was forbidden from sharing that data with the authors of this study during the course of this HuGL study, as is standard for blinded clinical trials. Any conclusions in this manuscript regarding relationships between HuGL mouse models and humans were made by the other authors, without input from W.R.S. related to any clinical trial activities.

### Competing interests

W.R.S. is an inventor on a patent application submitted by IAVI and The Scripps Research Institute that covers the eOD-GT8 immunogen.

## MATERIALS AND METHODS

### Generation of Mice

Paired HC and LC sequences were selected from a panel of cloned BCRs that were selected from individual healthy HIV^−^ human donors. B cells were selected for VRC01-class criteria, namely, coexpressing the VH1-2*02 allele and a LC with a 5 amino acid-long CDR3. All experimental mice were heterozygous for indicated HCs and LCs. All HuGL mice were on CD45.2 background while recipient mice were CD45.1 background. CD45.1 mice (B6.SJL-*Ptprc^a^Pepc^b^/BoyJ, stock No:002014)* were originally purchased from Jackson Labs (During breeding transgenic mice were maintained as heavy or light only. In some experiments donor heterozygous mice were generated from outbreeding to B6 females. HuGL mice used in experiments were bred by mating homozygous HC knock-in mice with corresponding homozygous LC knock-ins.

### HC knock-in mouse generation

Generation of HuGL 16, 17 and 18 HC mice was carried out essentially as described (*12, 47, 48*), by substituting the HuGL 16, 17 and 18 VDJ exon in the targeting construct by overlap PCR. Briefly, linearized targeting construct DNA was introduced into C57BL/6-derived embryonic stem cells and selected in media supplemented with G418. Cells that insert the gene non-specifically carry the DTA gene, which is toxic and cells lacking the neomycin resistance gene are counterselected. Positive clones were identified initially by a PCR strategy. The upstream promoter, leader and intron elements were from VHJ558.85.191. Targeting, embryonic stem cell screening, and mouse generation were as described (*12, 47, 48*).

### Light chain knock-in mouse generation

#### gRNAs design and Donor DNA plasmid preparation

gRNAs targeting the *IgKJ* region of the reference C57BL/6J genome (GRCm38) were designed using the GPP webtool (https://portals.broadinstitute.org/gpp/public/analysis-tools/sgrna-design). Selected gRNAs shown in Figure 1 B and C were synthesized by IDT and cloned into eSpCas9(1.1) (a gift from Feng Zhang Addgene plasmid # 71814). The LC donor DNA of HuGL18 was designed as follows from 5′ to 3′: a 2 kb homology arm upstream of the mouse *IgK-J5;* the mouse *IGKV4-53* LC promoter starting 1553-basepair upstream of, and including, the leader exon and intron; HuGL18 VJ; and a 1934-base region of DNA downstream of Jκ5 including the splicing signal. As for HuGL16 and 17 donor plasmids, the only difference in the backbone was the 5′ arm homology, which incuded a stretch of 2105 bp upstream of the IgKJ1-gRNA#2 targeting site. The homology regions and promoter were amplified from C57BL/6J genomic DNA. All LC fragments were synthesized (IDT). All fragments were assembled using the ligase-independent cloning method into pBKS (Addgene, 212207) and transformed into NEB® Stable Competent E. coli (NEB, C3040H). All plasmids were sequenced using several primers to generate high quality coverage of the entire insertions. Donor DNA plasmids were purified using EndoFree plasmid maxi kit (Qiagen, #12362) for engineering test and zygote microinjection.

#### gRNAs optimization in pro-B cells

*Rag1*^−/−^ pro-B cells were cultured in complete medium, RPMI- 1640 (Invitrogen, 21870076) supplemented with non-essential amino acids (Invitrogen,11140050), Sodium pyruvate (Invitrogen, 11360070), 55 μM 2-mercaptoethanol (Invitrogen, 21985023), 10% FBS (Invitrogen, 26140-079) and 100 units of Penicillin and 0.1 mg/ml of Streptomycin (P/S) (Invitrogen, 15140122). The cultured cells were washed twice with DPBS (Invitrogen, 14190-144) and resuspended at 0.5 million cells/ 10 μl in Neon R buffer (Invitrogen, MPK10096). 1 μg of each gRNA, 1 μg Donor DNA and 1 μg EGFP were mixed and added to the resuspended cells. The cells were electroporated using a 10 μl tip (Invitrogen, MPK1096) with 1400 V, 25 ms, 1 pulse using a Neon electroporation machine (Invitrogen). Transfected cells were immediately cultured in complete medium without P/S for 1 h before P/S was added for further culture. 48 hours later, the engineered cells were fixed with 4% PFA in DPBS at RT for 15 minutes and washed twice with FACS buffer (2% FBS in DPBS). Anti-mouse kappa LC antibody clone 187.1 was used to detect the engineering efficiency. EGFP-positive cells were gated for further engineering efficiency analysis.

#### Zygote injections

The gRNAs giving the highest engineering efficiency were chosen to generate LC knock-in mice using a Cas9 ribonucleoprotein approach along with the same donor plasmids described above. Female C57BL/6J mice (24-28 days old) were super-ovulated by the i.p. injection of 5 IU of pregnant mare serum gonadotropin (PMSG), followed by the 5 IU i.p. injection of human chorionic gonadotropin (hCG) 47 h later, then the mice were mated to C57BL/6J stud males. Fertilized oocytes were collected from the oviducts 21-22 h post hCG injection. Cas9 protein (50 ng/μl), sgRNA (5 ng/μl) and donor plasmid (10 ng/μl) were mixed in IDTE 1X TE nuclease-free buffer (pH 7.5) and injected into the pronuclei of fertilized eggs in a droplet of FHM medium containing 5 μg/ ml of cytochalasin B, using a FemtoJet microinjector (Eppendorf) with a constant flow setting. The injected zygotes were cultured in KSOM medium with amino acids at 37°C under 5% CO2 overnight to 2-cell stage and surgically transferred into oviducts of pseudopregnant CD-1 recipient females at 0.5 dpc.

### Flow cytometry analysis

Spleen samples were prepared as previously described (*62*). In brief, spleen suspensions were generated by smashing the spleen between frosted glass slides. Red blood cells were lysed with ammonium chloride (0.83%) before filtering cells through a 40 μM cell strainer to generate single-cell suspensions. Fc Blocker (homemade mAb 2G4) was added to single-cell suspensions at 1 μg per 10^6^ cells before antibody staining. For antigen-specific staining, tetramerized GT8 and GT8- KO11 were made by mixing biotinylated AviTagged GT8 and GT8-KO11 monomer with Streptavidin-AF647 and Streptavidin-AF488 respectively. Fluorophore-conjugated antibodies CD19 (Biolegend, 115540, 152408; BD Biosciences, 566107), CD45.1 (Biolegend, 110728, 110738), TCRb (Biolegend, 109228), GL7 (Biolegend, 144608, 144610), F4/80 (Biolegend, 123128), Ter119 (Biolegend, 116228), CD38 (Biolegend, 102718; BD Biosciences, 740245), CD45.2 (Biolegend, 109806, 109822), IgM (Biolegend, 314524), IgD (Biolegend, 405721), IgG1 (Biolegend, 406620, 406616), CD80 (Biolegend, 104712), CD73 (Biolegend, 127210), PD-L2 (Biolegend, 107216), CXCR4 (Biolegend, 146511), CD86 (Biolegend, 105008), PD1 (Biolegend, 135214), CD8 (Biolegend, 100722), CD4 (homemade AF-647), CXCR5 (Biolegend, 145512), BCL6 (Biolegend, 648304), B220 (Biolegend, 103236), CD21 () CD23 () were used to define different cellular populations. CD45.1 and CD45.2 mAbs were used to distinguish host and transferred HuGL^HL^ B cells respectively. IgG1 staining was used to identify class-switched B cells; Germinal center (GC) B cells were gated as CD19^+^TCRb^-^CD38^-^GL7^+^ and HuGL^HL^ fraction further gated as CD45.1^-^ CD45.2^+^; HuGL^HL^ Memory B cells (MCs) were gated as Live, CD45.1^-^ CD45.2^+^ CD19^+^ CD38^+^ TCRβ^-^ IgD^-^.

### Calcium flux analysis

Calcium flux was monitored as previously described with minor modification (*63*). Primary B cells from HuGL16^HL^, HuGL17^HL^, HuGL18^HL^, and WT mice were isolated using negative isolation kit (Miltenyi, 130-090-862). B cells were suspended at 4 million cells/ml in Advanced DMEM, labeled with 1 μM Indo-1 (Invitrogen, I1223) for 1 hour at 37°C, washed twice with DPBS, and resuspended in Advanced DMEM supplemented with 10% FBS. Basal Indo-1 fluorescence was recorded for 60 seconds, then antigens were added at 10 μg/ml final concentration and Ca^++^ signals were recorded for 180 s before ionomycin was added. Calcium flux analysis was performed on an LSRII cytometer (BD, San Jose, CA). Kinetic analysis was performed using FlowJo (Tree Star, Ashland, OR).

### Single-cell sorting and sequencing

Single cell sequencing was performed using one of two methods. Most sequencing was done using the RT-PCR based method. The results between the methods highly agreed and were considered equivalent.

Method 1: Single cell PCR for HuGL18 and HuGL17 was performed as in (*27*) which was adapted from Von Boehmer et al. HuGL18 HC sequencing primers were as previously reported (*27*). Briefly, cDNA was generated from RNA from single cell sorted IgG1^+^ GC B cells and then nested PCRs were performed to amplify both heavy and light chains. HuGL18 HCs and HuGL17 HCs were sequenced using 2MRG and 2MFG from Abbott et al 2018. HuGL18 LC sequencing primers were optimized and as follows VK320FM1 CATATTGTCCAGTGGAGAA and VK320RM3 CCGTTTGATTTCCACCTT. HuGL17 LC sequencing primers were as follows: VK15FM1 GACATCCAGATGACCCAG, VK15RM2 TTGATTTCCACCTTGGTCC. Sequences were aligned and trimmed in Sequencher v5.1. Analysis of VRC01-class mutations was performed as in (*27*).

Method 2: Class-switched IgG1^+^HuGL^HL^ GCs from spleen were individually sorted into wells of 96-well plate containing 10 μl of DPBS. Cells were spun down, frozen on dry ice and stored at −80 °C. To amplify HuGL heavy and LCs from single frozen cells, gDNA was prepared by adding 11 μl 2 X lysis buffer (20 mM Tris-HCl (pH 7.6), 100mM NaCl, 12.5 mM MgCl_2_, 0.09% Tween-20) containing 0.2 mg/ml protease K (NEB, P8107S) and incubating at 56°C for 1 hour. Protease was heat inactivated at 98°C for 15 min. The nested PCR is used for amplifying the HC and LC. For outer PCR, the 30 μl outer Phusion (Thermo, F530L) PCR master including both HC and LC outer sets of primers was directly added to the 96-well plate for amplifying HC and LC. 1 μl PCR products from outer PCR amplification was used for inner PCR amplification. The PCR products were subjected to electrophoresis through 1.5% agarose.

### B cell isolation and transfers

HuGL16, HuGL17, and HuGL18 B cells were all treated as in (*27*). Briefly spleens were collected, manually dissociated and filtered. Lymphocytes were separated by Ficoll gradient. Lymphocytes were then stained with anti-CD43 FITC (clone S7, BD Biosciences) for 30 minutes. Cells were washed, stained with anti-FITC microbeads (Miltenyi Biotech), washed, filtered and then passed through a LS Column on a Quadromacs magnet. EDTA was specifically avoided to maintain cell health. LS column was washed once with 2ml of buffer which was discarded prior to running cells. Collected cells were then checked by flow cytometry for B cell purity and antigen-positive cells using fluorescently labeled eOD-GT8. Cells were enumerated on a hemocytometer. Final tubes, syringes, and pipette tips were pre-coated for at least 1 hour with buffer. All steps were performed with 5% horse serum/DPBS (with Ca^++^ and Mg^++^).

### Immunogen Production

eOD-GT8 d41m3 was produced as previously described (*23*). Care was taken to evaluate immunogens for low endotoxin. The eOD-GT8-KO used was eOD-GT8-KO11 (*64*). Affinities for HuGL16, HuGL17, and HuGL18 for eOD-GT8 were previously published, and each were subsequently re-measured on multiple occasions, against different batches of eOD-GT8, and an representative K_d_ is shown.

### ELISAs

ELISAs for eOD-GT8 and eOD-GT8-KO were performed as in (*27*) with the only modification being the use of 96-well polysorb NUNC plates (Thermo). 60mers were used for all ELISAs at a coating concentration of 2 μg/ml. For negative control ELISA in Figure 2A (bottom) mice received a low number of HuGL18 B cells (1 in 10^6^ B cells).

### FACS and cell sorting

All flow cytometry analysis was in general performed as in (*27*). Briefly, all single cell suspensions were FC blocked for 10 minutes (clone 2.4G2) prior to all stains. Primary stains were carried out for 30 minutes. If no secondary stains were in a given panel, cells were washed twice with 10x volume (1 ml FACS buffer for 100 μL staining buffer) before fixation or collection on cytometer. 3-laser 12-color FACSCelesta was used for most experiments. Some experiments were collected while sorting on Aria Fusion. Data was processed using Flowjo 10. All mAb clones used were as listed in (*27*).

For sorting, 4-way purity mode was used and single cells were sorted into 96 well plate with collection buffer. Collection buffer and plates were the same as reported in Abbott et al 2018. Single cell mode was avoided as it was too restrictive. Cells were sorted with the 70 μ nozzle with custom pressure (61.5PSI) and sort rate was kept at 2.4 or below to ensure targeting. ACDU was checked for targeting after each plate and adjusted if necessary. Cells were kept at 4C (not on ice) while awaiting sort to avoid cell death. No in-line filter was used during these sorts.

### Histology

Histology was done essentially as in (*27*). Briefly 5-8 μM frozen splenic sections were cut, fixed in 1:1 mixture of acetone/methanol at −20C for 10 minutes and dried. Upon rehydration, slides were blocked (FC block and 0.5% BSA in PBS) and then stained with indicated antibodies. Zeiss axioscan was used for acquisition. Zen software was used for evaluation images.

## REFERENCES

1. D. E. Bloom, V. Y. Fan, J. P. Sevilla, The broad socioeconomic benefits of vaccination. Sci Transl Med 10, (2018).

2. S. A. Plotkin, W. A. Orenstein, P. A. Offit, Plotkin’s vaccines. (Elsevier, Philadelphia, PA, ed. Seventh edition., 2018), pp. xxi, 1691 pages.

3. C. Havenar-Daughton, J. H. Lee, S. Crotty, Tfh cells and HIV bnAbs, an immunodominance model of the HIV neutralizing antibody generation problem. Immunol. Rev. 275, 49–61 (2017).

4. P. Piot, H. J. Larson, K. L. O’Brien, J. N’Kengasong, E. Ng, S. Sow, B. Kampmann, Immunization: vital progress, unfinished agenda. Nature 575, 119–129 (2019).

5. P. D. Kwong, J. R. Mascola, HIV-1 Vaccines Based on Antibody Identification, B Cell Ontogeny, and Epitope Structure. Immunity 48, 855–871 (2018).

6. D. R. Burton, What Are the Most Powerful Immunogen Design Vaccine Strategies? Reverse Vaccinology 2.0 Shows Great Promise. Cold Spring Harb Perspect Biol 9, (2017).

7. R. Rappuoli, M. J. Bottomley, U. D’Oro, O. Finco, E. De Gregorio, Reverse vaccinology 2.0: Human immunology instructs vaccine antigen design. J. Exp. Med. 213, 469–481 (2016).

8. J. Jardine, J. P. Julien, S. Menis, T. Ota, O. Kalyuzhniy, A. McGuire, D. Sok, P. S. Huang, S. Macpherson, M. Jones, T. Nieusma, J. Mathison, D. Baker, A. B. Ward, D. R. Burton, L. Stamatatos, D. Nemazee, I. A. Wilson, W. R. Schief, Rational HIV Immunogen Design to Target Specific Germline B Cell Receptors. Science 340, 711–716 (2013).

9. J. G. Jardine, D. Sok, J. P. Julien, B. Briney, A. Sarkar, C. H. Liang, E. A. Scherer, C. J. Henry Dunand, Y. Adachi, D. Diwanji, J. Hsueh, M. Jones, O. Kalyuzhniy, M. Kubitz, S. Spencer, M. Pauthner, K. L. Saye-Francisco, F. Sesterhenn, P. C. Wilson, D. M. Galloway, R. L. Stanfield, I. A. Wilson, D. R. Burton, W. R. Schief, Minimally Mutated HIV-1 Broadly Neutralizing Antibodies to Guide Reductionist Vaccine Design. PLoS Pathog 12, e1005815 (2016).

10. E. Landais, P. L. Moore, Development of broadly neutralizing antibodies in HIV-1 infected elite neutralizers. Retrovirology 15, 61 (2018).

11. P. Dosenovic, L. von Boehmer, A. Escolano, J. Jardine, N. T. Freund, A. D. Gitlin, A. T. McGuire, D. W. Kulp, T. Oliveira, L. Scharf, J. Pietzsch, M. D. Gray, A. Cupo, M. J. van Gils, K. H. Yao, C. Liu, A. Gazumyan, M. S. Seaman, P. J. Bjorkman, R. W. Sanders, J. P. Moore, L. Stamatatos, W. R. Schief, M. C. Nussenzweig, Immunization for HIV-1 Broadly Neutralizing Antibodies in Human Ig Knockin Mice. Cell 161, 1505–1515 (2015).

12. J. G. Jardine, T. Ota, D. Sok, M. Pauthner, D. W. Kulp, O. Kalyuzhniy, P. D. Skog, T. C. Thinnes, D. Bhullar, B. Briney, S. Menis, M. Jones, M. Kubitz, S. Spencer, Y. Adachi, D. R. Burton, W. R. Schief, D. Nemazee, HIV-1 VACCINES. Priming a broadly neutralizing antibody response to HIV-1 using a germline-targeting immunogen. Science 349, 156–161 (2015).

13. M. Medina-Ramirez, F. Garces, A. Escolano, P. Skog, S. W. de Taeye, I. Del Moral-Sanchez, A. T. McGuire, A. Yasmeen, A. J. Behrens, G. Ozorowski, T. van den Kerkhof, N. T. Freund, P. Dosenovic, Y. Hua, A. D. Gitlin, A. Cupo, P. van der Woude, M. Golabek, K. Sliepen, T. Blane, N. Kootstra, M. J. van Breemen, L. K. Pritchard, R. L. Stanfield, M. Crispin, A. B. Ward, L. Stamatatos, P. J. Klasse, J. P. Moore, D. Nemazee, M. C. Nussenzweig, I. A. Wilson, R. W. Sanders, Design and crystal structure of a native-like HIV-1 envelope trimer that engages multiple broadly neutralizing antibody precursors in vivo. J. Exp. Med. 214, 2573–2590 (2017).

14. J. M. Steichen, D. W. Kulp, T. Tokatlian, A. Escolano, P. Dosenovic, R. L. Stanfield, L. E. McCoy, G. Ozorowski, X. Hu, O. Kalyuzhniy, B. Briney, T. Schiffner, F. Garces, N. T. Freund, A. D. Gitlin, S. Menis, E. Georgeson, M. Kubitz, Y. Adachi, M. Jones, A. A. Mutafyan, D. S. Yun, C. T. Mayer, A. B. Ward, D. R. Burton, I. A. Wilson, D. J. Irvine, M. C. Nussenzweig, W. R. Schief, HIV Vaccine Design to Target Germline Precursors of Glycan-Dependent Broadly Neutralizing Antibodies. Immunity 45, 483–496 (2016).

15. A. Escolano, H. B. Gristick, M. E. Abernathy, J. Merkenschlager, R. Gautam, T. Y. Oliveira, J. Pai, A. P. West, Jr., C. O. Barnes, A. A. Cohen, H. Wang, J. Golijanin, D. Yost, J. R. Keeffe, Z. Wang, P. Zhao, K. H. Yao, J. Bauer, L. Nogueira, H. Gao, A. V. Voll, D. C. Montefiori, M. S. Seaman, A. Gazumyan, M. Silva, A. T. McGuire, L. Stamatatos, D. J. Irvine, L. Wells, M. A. Martin, P. J. Bjorkman, M. C. Nussenzweig, Immunization expands B cells specific to HIV-1 V3 glycan in mice and macaques. Nature 570, 468–473 (2019).

16. C. C. LaBranche, R. Henderson, A. Hsu, S. Behrens, X. Chen, T. Zhou, K. Wiehe, K. O. Saunders, S. M. Alam, M. Bonsignori, M. J. Borgnia, Q. J. Sattentau, A. Eaton, K. Greene, H. Gao, H. X. Liao, W. B. Williams, J. Peacock, H. Tang, L. G. Perez, R. J. Edwards, T. B. Kepler, B. T. Korber, P. D. Kwong, J. R. Mascola, P. Acharya, B. F. Haynes, D. C. Montefiori, Neutralization-guided design of HIV-1 envelope trimers with high affinity for the unmutated common ancestor of CH235 lineage CD4bs broadly neutralizing antibodies. PLoS Pathog 15, e1008026 (2019).

17. K. R. Parks, A. J. MacCamy, J. Trichka, M. Gray, C. Weidle, A. J. Borst, A. Khechaduri, B. Takushi, P. Agrawal, J. Guenaga, R. T. Wyatt, R. Coler, M. Seaman, C. LaBranche, D. C. Montefiori, D. Veesler, M. Pancera, A. McGuire, L. Stamatatos, Overcoming Steric Restrictions of VRC01 HIV-1 Neutralizing Antibodies through Immunization. Cell Rep 29, 3060–3072 e3067 (2019).

18. A. T. McGuire, S. Hoot, A. M. Dreyer, A. Lippy, A. Stuart, K. W. Cohen, J. Jardine, S. Menis, J. F. Scheid, A. P. West, W. R. Schief, L. Stamatatos, Engineering HIV envelope protein to activate germline B cell receptors of broadly neutralizing anti-CD4 binding site antibodies. J. Exp. Med. 210, 655–663 (2013).

19. M. Bonsignori, E. Scott, K. Wiehe, D. Easterhoff, S. M. Alam, K. K. Hwang, M. Cooper, S. M. Xia, R. Zhang, D. C. Montefiori, R. Henderson, X. Nie, G. Kelsoe, M. A. Moody, X. Chen, M. G. Joyce, P. D. Kwong, M. Connors, J. R. Mascola, A. T. McGuire, L. Stamatatos, M. Medina-Ramirez, R. W. Sanders, K. O. Saunders, T. B. Kepler, B. F. Haynes, Inference of the HIV-1 VRC01 Antibody Lineage Unmutated Common Ancestor Reveals Alternative Pathways to Overcome a Key Glycan Barrier. Immunity, (2018).

20. J. P. Julien, D. Sok, R. Khayat, J. H. Lee, K. J. Doores, L. M. Walker, A. Ramos, D. C. Diwanji, R. Pejchal, A. Cupo, U. Katpally, R. S. Depetris, R. L. Stanfield, R. McBride, A. J. Marozsan, J. C. Paulson, R. W. Sanders, J. P. Moore, D. R. Burton, P. Poignard, A. B. Ward, I. A. Wilson, Broadly neutralizing antibody PGT121 allosterically modulates CD4 binding via recognition of the HIV-1 gp120 V3 base and multiple surrounding glycans. PLoS Pathog 9, e1003342 (2013).

21. H. Mouquet, L. Scharf, Z. Euler, Y. Liu, C. Eden, J. F. Scheid, A. Halper-Stromberg, P. N. Gnanapragasam, D. I. Spencer, M. S. Seaman, H. Schuitemaker, T. Feizi, M. C. Nussenzweig, P. J. Bjorkman, Complex-type N-glycan recognition by potent broadly neutralizing HIV antibodies. Proc Natl Acad Sci U S A 109, E3268–3277 (2012).

22. M. Tian, C. Cheng, X. Chen, H. Duan, H. L. Cheng, M. Dao, Z. Sheng, M. Kimble, L. Wang, S. Lin, S. D. Schmidt, Z. Du, M. G. Joyce, Y. Chen, B. J. DeKosky, Y. Chen, E. Normandin, E. Cantor, R. E. Chen, N. A. Doria-Rose, Y. Zhang, W. Shi, W. P. Kong, M. Choe, A. R. Henry, F. Laboune, I. S. Georgiev, P. Y. Huang, S. Jain, A. T. McGuire, E. Georgeson, S. Menis, D. C. Douek, W. R. Schief, L. Stamatatos, P. D. Kwong, L. Shapiro, B. F. Haynes, J. R. Mascola, F. W. Alt, Induction of HIV Neutralizing Antibody Lineages in Mice with Diverse Precursor Repertoires. Cell 166, 1471–1484 e1418 (2016).

23. D. Sok, B. Briney, J. G. Jardine, D. W. Kulp, S. Menis, M. Pauthner, A. Wood, E. C. Lee, K. M. Le, M. Jones, A. Ramos, O. Kalyuzhniy, Y. Adachi, M. Kubitz, S. MacPherson, A. Bradley, G. A. Friedrich, W. R. Schief, D. R. Burton, Priming HIV-1 broadly neutralizing antibody precursors in human Ig loci transgenic mice. Science, (2016).

24. P. Dosenovic, E. E. Kara, A. K. Pettersson, A. T. McGuire, M. Gray, H. Hartweger, E. S. Thientosapol, L. Stamatatos, M. C. Nussenzweig, Anti-HIV-1 B cell responses are dependent on B cell precursor frequency and antigen-binding affinity. Proc Natl Acad Sci U S A 115, 4743–4748 (2018).

25. A. Escolano, J. M. Steichen, P. Dosenovic, D. W. Kulp, J. Golijanin, D. Sok, N. T. Freund, A. D. Gitlin, T. Oliveira, T. Araki, S. Lowe, S. T. Chen, J. Heinemann, K. H. Yao, E. Georgeson, K. L. Saye-Francisco, A. Gazumyan, Y. Adachi, M. Kubitz, D. R. Burton, W. R. Schief, M. C. Nussenzweig, Sequential Immunization Elicits Broadly Neutralizing Anti-HIV-1 Antibodies in Ig Knockin Mice. Cell 166, 1445–1458 e1412 (2016).

26. A. T. McGuire, M. D. Gray, P. Dosenovic, A. D. Gitlin, N. T. Freund, J. Petersen, C. Correnti, W. Johnsen, R. Kegel, A. B. Stuart, J. Glenn, M. S. Seaman, W. R. Schief, R. K. Strong, M. C. Nussenzweig, L. Stamatatos, Specifically modified Env immunogens activate B-cell precursors of broadly neutralizing HIV-1 antibodies in transgenic mice. Nat Commun 7, 10618 (2016).

27. R. K. Abbott, J. H. Lee, S. Menis, P. Skog, M. Rossi, T. Ota, D. W. Kulp, D. Bhullar, O. Kalyuzhniy, C. Havenar-Daughton, W. R. Schief, D. Nemazee, S. Crotty, Precursor Frequency and Affinity Determine B Cell Competitive Fitness in Germinal Centers, Tested with Germline-Targeting HIV Vaccine Immunogens. Immunity 48, 133–146 e136 (2018).

28. J. G. Jardine, D. W. Kulp, C. Havenar-Daughton, A. Sarkar, B. Briney, D. Sok, F. Sesterhenn, J. Ereno-Orbea, O. Kalyuzhniy, I. Deresa, X. Hu, S. Spencer, M. Jones, E. Georgeson, Y. Adachi, M. Kubitz, A. C. deCamp, J. P. Julien, I. A. Wilson, D. R. Burton, S. Crotty, W. R. Schief, HIV-1 broadly neutralizing antibody precursor B cells revealed by germline-targeting immunogen. Science 351, 1458–1463 (2016).

29. J. M. Steichen, Y. C. Lin, C. Havenar-Daughton, S. Pecetta, G. Ozorowski, J. R. Willis, L. Toy, D. Sok, A. Liguori, S. Kratochvil, J. L. Torres, O. Kalyuzhniy, E. Melzi, D. W. Kulp, S. Raemisch, X. Hu, S. M. Bernard, E. Georgeson, N. Phelps, Y. Adachi, M. Kubitz, E. Landais, J. Umotoy, A. Robinson, B. Briney, I. A. Wilson, D. R. Burton, A. B. Ward, S. Crotty, F. D. Batista, W. R. Schief, A generalized HIV vaccine design strategy for priming of broadly neutralizing antibody responses. Science 366, (2019).

30. A Phase I Trial to Evaluate the Safety and Immunogenicity of eOD-GT8 60mer Vaccine, Adjuvanted. NCT03547245 G001.

31. T. Zhou, I. Georgiev, X. Wu, Z. Y. Yang, K. Dai, A. Finzi, Y. D. Kwon, J. F. Scheid, W. Shi, L. Xu, Y. Yang, J. Zhu, M. C. Nussenzweig, J. Sodroski, L. Shapiro, G. J. Nabel, J. R. Mascola, P. D. Kwong, Structural basis for broad and potent neutralization of HIV-1 by antibody VRC01. Science 329, 811–817 (2010).

32. T. Zhou, J. Zhu, X. Wu, S. Moquin, B. Zhang, P. Acharya, I. S. Georgiev, H. R. Altae-Tran, G. Y. Chuang, M. G. Joyce, Y. Do Kwon, N. S. Longo, M. K. Louder, T. Luongo, K. McKee, C. A. Schramm, J. Skinner, Y. Yang, Z. Yang, Z. Zhang, A. Zheng, M. Bonsignori, B. F. Haynes, J. F. Scheid, M. C. Nussenzweig, M. Simek, D. R. Burton, W. C. Koff, N. C. S. Program, J. C. Mullikin, M. Connors, L. Shapiro, G. J. Nabel, J. R. Mascola, P. D. Kwong, Multidonor analysis reveals structural elements, genetic determinants, and maturation pathway for HIV-1 neutralization by VRC01-class antibodies. Immunity 39, 245–258 (2013).

33. J. Huang, B. H. Kang, E. Ishida, T. Zhou, T. Griesman, Z. Sheng, F. Wu, N. A. Doria-Rose, B. Zhang, K. McKee, S. O’Dell, G. Y. Chuang, A. Druz, I. S. Georgiev, C. A. Schramm, A. Zheng, M. G. Joyce, M. Asokan, A. Ransier, S. Darko, S. A. Migueles, R. T. Bailer, M. K. Louder, S. M. Alam, R. Parks, G. Kelsoe, T. Von Holle, B. F. Haynes, D. C. Douek, V. Hirsch, M. S. Seaman, L. Shapiro, J. R. Mascola, P. D. Kwong, M. Connors, Identification of a CD4-Binding-Site Antibody to HIV that Evolved Near-Pan Neutralization Breadth. Immunity 45, 1108–1121 (2016).

34. J. Umotoy, B. S. Bagaya, C. Joyce, T. Schiffner, S. Menis, K. L. Saye-Francisco, T. Biddle, S. Mohan, T. Vollbrecht, O. Kalyuzhniy, S. Madzorera, D. Kitchin, B. Lambson, M. Nonyane, W. Kilembe, I. P. C. Investigators, I. A. H. R. Network, P. Poignard, W. R. Schief, D. R. Burton, B. Murrell, P. L. Moore, B. Briney, D. Sok, E. Landais, Rapid and Focused Maturation of a VRC01-Class HIV Broadly Neutralizing Antibody Lineage Involves Both Binding and Accommodation of the N276-Glycan. Immunity 51, 141–154 e146 (2019).

35. M. M. Sajadi, A. Dashti, Z. Rikhtegaran Tehrani, W. D. Tolbert, M. S. Seaman, X. Ouyang, N. Gohain, M. Pazgier, D. Kim, G. Cavet, J. Yared, R. R. Redfield, G. K. Lewis, A. L. DeVico, Identification of Near-Pan-neutralizing Antibodies against HIV-1 by Deconvolution of Plasma Humoral Responses. Cell 173, 1783–1795 e1714 (2018).

36. X. Wu, Z. Y. Yang, Y. Li, C. M. Hogerkorp, W. R. Schief, M. S. Seaman, T. Zhou, S. D. Schmidt, L. Wu, L. Xu, N. S. Longo, K. McKee, S. O’Dell, M. K. Louder, D. L. Wycuff, Y. Feng, M. Nason, N. Doria-Rose, M. Connors, P. D. Kwong, M. Roederer, R. T. Wyatt, G. J. Nabel, J. R. Mascola, Rational design of envelope identifies broadly neutralizing human monoclonal antibodies to HIV-1. Science 329, 856–861 (2010).

37. G. D. Victora, M. C. Nussenzweig, Germinal centers. Annu. Rev. Immunol. 30, 429–457 (2012).

38. J. G. Cyster, C. D. C. Allen, B Cell Responses: Cell Interaction Dynamics and Decisions. Cell 177, 524–540 (2019).

39. S. Crotty, T Follicular Helper Cell Biology: A Decade of Discovery and Diseases. Immunity 50, 1132–1148 (2019).

40. L. M. Corcoran, D. M. Tarlinton, Regulation of germinal center responses, memory B cells and plasma cell formation-an update. Curr. Opin. Immunol. 39, 59–67 (2016).

41. O. Bannard, J. G. Cyster, Germinal centers: programmed for affinity maturation and antibody diversification. Curr. Opin. Immunol. 45, 21–30 (2017).

42. G. D. Victora, H. Mouquet, What Are the Primary Limitations in B-Cell Affinity Maturation, and How Much Affinity Maturation Can We Drive with Vaccination? Lessons from the Antibody Response to HIV-1. Cold Spring Harb Perspect Biol 10, (2018).

43. C. Havenar-Daughton, A. Sarkar, D. W. Kulp, L. Toy, X. Hu, I. Deresa, O. Kalyuzhniy, K. Kaushik, A. A. Upadhyay, S. Menis, E. Landais, L. Cao, J. K. Diedrich, S. Kumar, T. Schiffner, S. M. Reiss, G. Seumois, J. R. Yates, J. C. Paulson, S. E. Bosinger, I. A. Wilson, W. R. Schief, S. Crotty, The human naive B cell repertoire contains distinct subclasses for a germline-targeting HIV-1 vaccine immunogen. Sci Transl Med 10, (2018).

44. H. Duan, X. Chen, J. C. Boyington, C. Cheng, Y. Zhang, A. J. Jafari, T. Stephens, Y. Tsybovsky, O. Kalyuzhniy, P. Zhao, S. Menis, M. C. Nason, E. Normandin, M. Mukhamedova, B. J. DeKosky, L. Wells, W. R. Schief, M. Tian, F. W. Alt, P. D. Kwong, J. R. Mascola, Glycan Masking Focuses Immune Responses to the HIV-1 CD4-Binding Site and Enhances Elicitation of VRC01-Class Precursor Antibodies. Immunity 49, 301–311 e305 (2018).

45. C. Havenar-Daughton, R. K. Abbott, W. R. Schief, S. Crotty, When designing vaccines, consider the starting material: the human B cell repertoire. Curr. Opin. Immunol. 53, 209–216 (2018).

46. B. Briney, D. Sok, J. G. Jardine, D. W. Kulp, P. Skog, S. Menis, R. Jacak, O. Kalyuzhniy, N. de Val, F. Sesterhenn, K. M. Le, A. Ramos, M. Jones, K. L. Saye-Francisco, T. R. Blane, S. Spencer, E. Georgeson, X. Hu, G. Ozorowski, Y. Adachi, M. Kubitz, A. Sarkar, I. A. Wilson, A. B. Ward, D. Nemazee, D. R. Burton, W. R. Schief, Tailored Immunogens Direct Affinity Maturation toward HIV Neutralizing Antibodies. Cell 166, 1459–1470 e1411 (2016).

47. C. Doyle-Cooper, K. E. Hudson, A. B. Cooper, T. Ota, P. Skog, P. E. Dawson, M. B. Zwick, W. R. Schief, D. R. Burton, D. Nemazee, Immune tolerance negatively regulates B cells in knock-in mice expressing broadly neutralizing HIV antibody 4E10. J. Immunol. 191, 3186–3191 (2013).

48. T. Ota, C. Doyle-Cooper, A. B. Cooper, K. J. Doores, M. Aoki-Ota, K. Le, W. R. Schief, R. T. Wyatt, D. R. Burton, D. Nemazee, B cells from knock-in mice expressing broadly neutralizing HIV antibody b12 carry an innocuous B cell receptor responsive to HIV vaccine candidates. J. Immunol. 191, 3179–3185 (2013).

49. K. O. Saunders, K. Wiehe, M. Tian, P. Acharya, T. Bradley, S. M. Alam, E. P. Go, R. Scearce, L. Sutherland, R. Henderson, A. L. Hsu, M. J. Borgnia, H. Chen, X. Lu, N. R. Wu, B. Watts, C. Jiang, D. Easterhoff, H. L. Cheng, K. McGovern, P. Waddicor, A. Chapdelaine-Williams, A. Eaton, J. Zhang, W. Rountree, L. Verkoczy, M. Tomai, M. G. Lewis, H. R. Desaire, R. J. Edwards, D. W. Cain, M. Bonsignori, D. Montefiori, F. W. Alt, B. F. Haynes, Targeted selection of HIV-specific antibody mutations by engineering B cell maturation. Science 366, (2019).

50. L. D’Souza, S. L. Gupta, V. Bal, S. Rath, A. George, CD73 expression identifies a subset of IgM(+) antigen-experienced cells with memory attributes that is T cell and CD40 signalling dependent. Immunology 152, 602–612 (2017).

51. F. Weisel, M. Shlomchik, Memory B Cells of Mice and Humans. Annu. Rev. Immunol. 35, 255–284 (2017).

52. D. Angeletti, J. W. Yewdell, Understanding and Manipulating Viral Immunity: Antibody Immunodominance Enters Center Stage. Trends Immunol. 39, 549–561 (2018).

53. M. Sangesland, L. Ronsard, S. W. Kazer, J. Bals, S. Boyoglu-Barnum, A. S. Yousif, R. Barnes, J. Feldman, M. Quirindongo-Crespo, P. M. McTamney, D. Rohrer, N. Lonberg, B. Chackerian, B. R. Graham, M. Kanekiyo, A. K. Shalek, D. Lingwood, Germline-Encoded Affinity for Cognate Antigen Enables Vaccine Amplification of a Human Broadly Neutralizing Response against Influenza Virus. Immunity 51, 735–749 e738 (2019).

54. M. Kuraoka, A. G. Schmidt, T. Nojima, F. Feng, A. Watanabe, D. Kitamura, S. C. Harrison, T. B. Kepler, G. Kelsoe, Complex Antigens Drive Permissive Clonal Selection in Germinal Centers. Immunity 44, 542–552 (2016).

55. D. Angeletti, J. S. Gibbs, M. Angel, I. Kosik, H. D. Hickman, G. M. Frank, S. R. Das, A. K. Wheatley, M. Prabhakaran, D. J. Leggat, A. B. McDermott, J. W. Yewdell, Defining B cell immunodominance to viruses. Nat. Immunol. 18, 456–463 (2017).

56. G. D. Victora, P. C. Wilson, Germinal center selection and the antibody response to influenza. Cell 163, 545–548 (2015).

57. L. Verkoczy, Y. Chen, H. Bouton-Verville, J. Zhang, M. Diaz, J. Hutchinson, Y. B. Ouyang, S. M. Alam, T. M. Holl, K. K. Hwang, G. Kelsoe, B. F. Haynes, Rescue of HIV-1 broad neutralizing antibody-expressing B cells in 2F5 VH x VL knockin mice reveals multiple tolerance controls. J. Immunol. 187, 3785–3797 (2011).

58. L. Verkoczy, Y. Chen, J. Zhang, H. Bouton-Verville, A. Newman, B. Lockwood, R. M. Scearce, D. C. Montefiori, S. M. Dennison, S. M. Xia, K. K. Hwang, H. X. Liao, S. M. Alam, B. F. Haynes, Induction of HIV-1 broad neutralizing antibodies in 2F5 knock-in mice: selection against membrane proximal external region-associated autoreactivity limits T-dependent responses. J. Immunol. 191, 2538–2550 (2013).

59. Y. C. Lin, S. Pecetta, J. M. Steichen, S. Kratochvil, E. Melzi, J. Arnold, S. K. Dougan, L. Wu, K. H. Kirsch, U. Nair, W. R. Schief, F. D. Batista, One-step CRISPR/Cas9 method for the rapid generation of human antibody heavy chain knock-in mice. EMBO J. 37, (2018).

60. J. T. Jacobsen, L. Mesin, S. Markoulaki, A. Schiepers, C. B. Cavazzoni, D. Bousbaine, R. Jaenisch, G. D. Victora, One-step generation of monoclonal B cell receptor mice capable of isotype switching and somatic hypermutation. J. Exp. Med. 215, 2686–2695 (2018).

61. L. Kong, B. Ju, Y. Chen, L. He, L. Ren, J. Liu, K. Hong, B. Su, Z. Wang, G. Ozorowski, X. Ji, Y. Hua, Y. Chen, M. C. Deller, Y. Hao, Y. Feng, F. Garces, R. Wilson, K. Dai, S. O’Dell, K. McKee, J. R. Mascola, A. B. Ward, R. T. Wyatt, Y. Li, I. A. Wilson, J. Zhu, Y. Shao, Key gp120 Glycans Pose Roadblocks to the Rapid Development of VRC01-Class Antibodies in an HIV-1- Infected Chinese Donor. Immunity 44, 939–950 (2016).

62. A. L. Gavin, D. Huang, C. Huber, A. Martensson, V. Tardif, P. D. Skog, T. R. Blane, T. C. Thinnes, K. Osborn, H. S. Chong, F. Kargaran, P. Kimm, A. Zeitjian, R. L. Sielski, M. Briggs, S. R. Schulz, A. Zarpellon, B. Cravatt, E. S. Pang, J. Teijaro, J. C. de la Torre, M. O’Keeffe, H. Hochrein, M. Damme, L. Teyton, B. R. Lawson, D. Nemazee, PLD3 and PLD4 are single-stranded acid exonucleases that regulate endosomal nucleic-acid sensing. Nat. Immunol. 19, 942–953 (2018).

63. T. Ota, C. Doyle-Cooper, A. B. Cooper, M. Huber, E. Falkowska, K. J. Doores, L. Hangartner, K. Le, D. Sok, J. Jardine, J. Lifson, X. Wu, J. R. Mascola, P. Poignard, J. M. Binley, B. K. Chakrabarti, W. R. Schief, R. T. Wyatt, D. R. Burton, D. Nemazee, Anti-HIV B Cell Lines as Candidate Vaccine Biosensors. J. Immunol. 189, 4816–4824 (2012).

64. M. Melo, E. Porter, Y. Zhang, M. Silva, N. Li, B. Dobosh, A. Liguori, P. Skog, E. Landais, S. Menis, D. Sok, D. Nemazee, W. R. Schief, R. Weiss, D. J. Irvine, Immunogenicity of RNA Replicons Encoding HIV Env Immunogens Designed for Self-Assembly into Nanoparticles. Mol. Ther., (2019).

