## Supplemental Figures 1-5 and Supplemental Table 1 for "B cells expressing authentic naive human VRC01-class BCRs can be primed and recruited to germinal centers in multiple independent mouse models"

* These authors contributed equally.

This PDF File includes

Fig S1: Analysis of pre-immune knock-in HuGL mice.

Fig S2: HuGL B cell calcium flux data.

Fig S3: HuGL18 day 16 extended mutation data.

Fig S4: HuGL 17 memory B cell data.

Fig S5: HuGL17 day 16 extended mutation data.

Table S1: Sequence features of HuGL B cells.

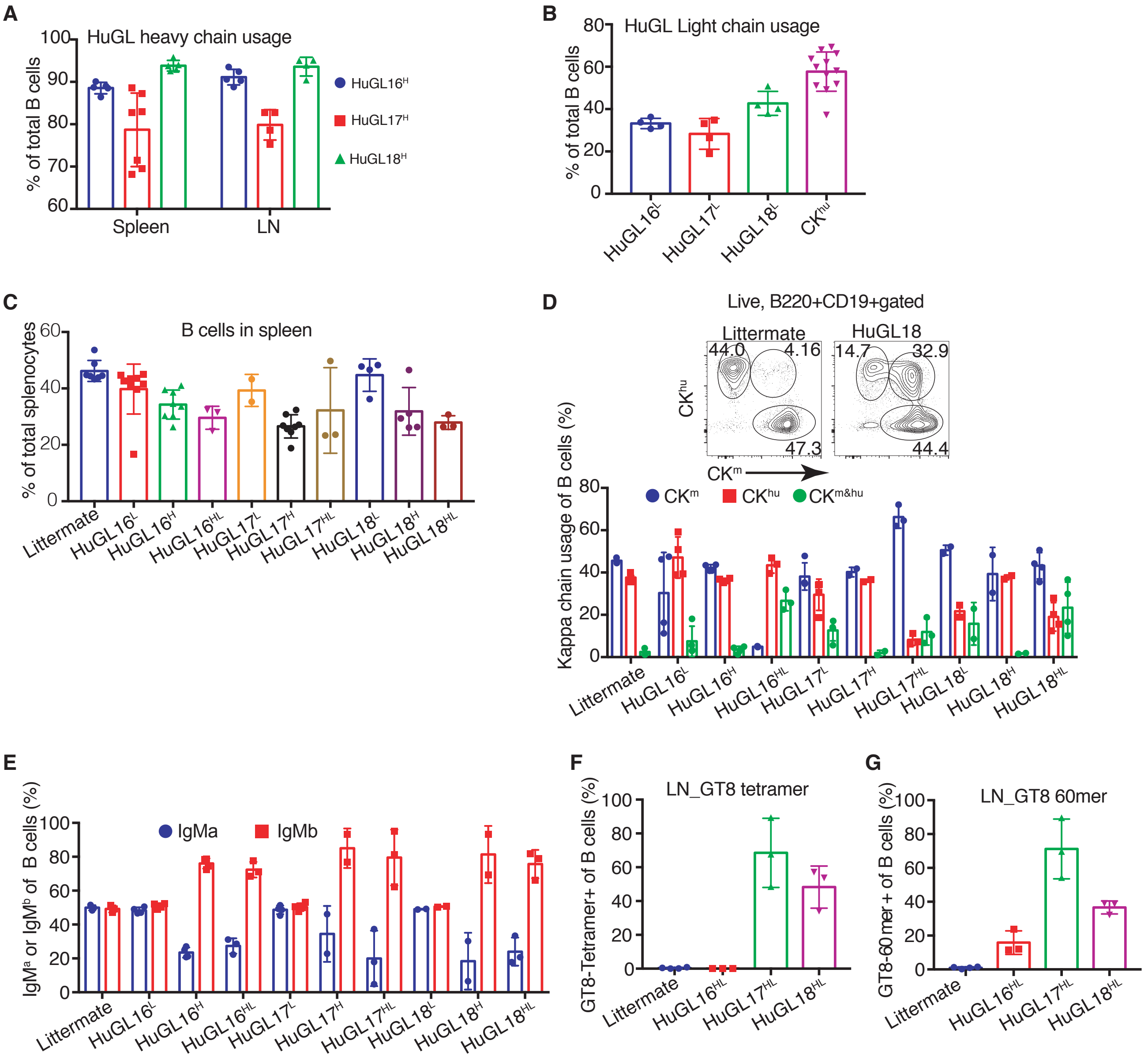
A
HuGL heavy chain usage
% of total B cells
Spleen
LN
HuGL16H
HuGL17H
HuGL18H
B
HuGL Light chain usage
% of total B cells
HuGL16L
HuGL17L
HuGL18L
CKhu
C
B cells in spleen
% of total splenocytes
Littermate
HuGL16L
HuGL16H
HuGL16HL
HuGL17L
HuGL17H
HuGL17HL
HuGL18L
HuGL18H
HuGL18HL
D
Live, B220+CD19+gated
Littermate
HuGL18
44.0
4.16
47.3
14.7
32.9
44.4
CKhu
CKm
Kappa chain usage of B cells (%)
CKm
CKhu
CKm&hu
E
IgMa or IgMb of B cells (%)
IgMa
IgMb
F
LN_GT8 tetramer
GT8-Tetramer+ of B cells (%)
G
LN_GT8 60mer
GT8-60 mer + of B cells (%)

Figure S1

**Figure S1.** Analysis of preimmune knock-in HuGL mice. The indicated BCR knock-in strains generated on the C57BL/6 background ($Igh^{b/b}C\kappa^{m/m}$) were bred to $IgH^{a/a}C\kappa^{hu/hu}$ mice and analyzed by flow cytometry for allelic exclusion and B cell numbers. (A) Analysis of HC allelic exclusion in the indicated HC-only mice. Spleen and lymph node B cells were assessed for expression of $IgM^b$ by flow cytometry. **(B)** LC-only strain mice were similarly assessed for expression of the knock-in allele. **(C)** Enumeration of B cell numbers as a percentage of spleen cells. **(D)** LC allelic exclusion analysis of H, L and H/L strains. Flow plot shows gating enumerating cells expressing knock-in vs endogenous κ allele (horizontal vs vertical axis, respectively); double positive cells are in the upper right quadrant. **(E)** IgH allelic exclusion of H, L and H/L bone marrow B cells. **(F,G)** Enumeration of GT8-binding cells quantitated with streptavidin tetramers **(F)** or 60mer nanoparticles **(G).**

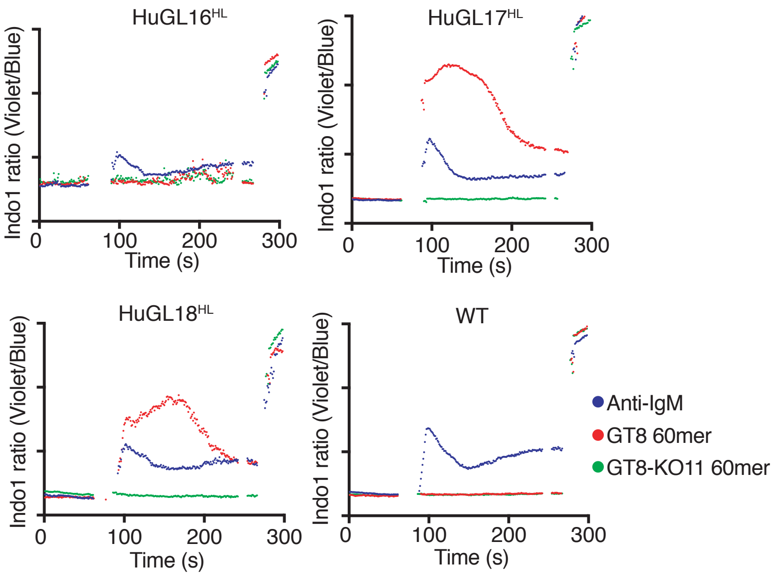

HuGL16^HL
HuGL17^HL
HuGL18^HL
WT
Indo1 ratio (Violet/Blue)
Time (s)
0
100
200
300
Anti-IgM
GT8 60mer
GT8-KO11 60mer


Figure S2

**Figure S2**. Calcium flux analysis of splenic B cells from HuGL16$^{HL}$, HuGL17$^{HL}$ and HuGL18$^{HL}$ mice treated with eOD-GT8 60mer nanoparticles, eOD-GT8-KO 60mer nanoparticles, or anti-IgM. Stimuli were applied at 10 μg/ml after 90 s equilibration followed by treatment with ionomycin to assess Indo1 loading.

**A**

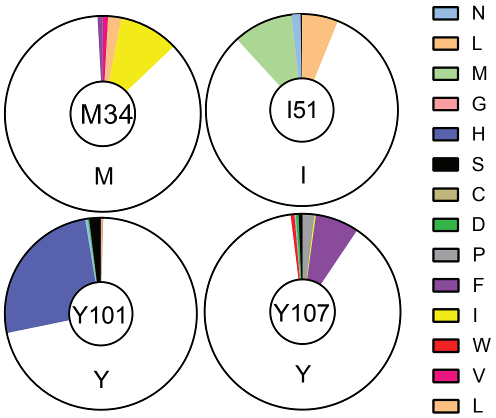


**Figure S3**

**Figure S3**. HuGL18 day 16 extended mutation data. Amino acid replacements in VH1-2 heavy chain are shown for HuGL18 GC B cells sorted from day 16 mice, as per Figure 5.

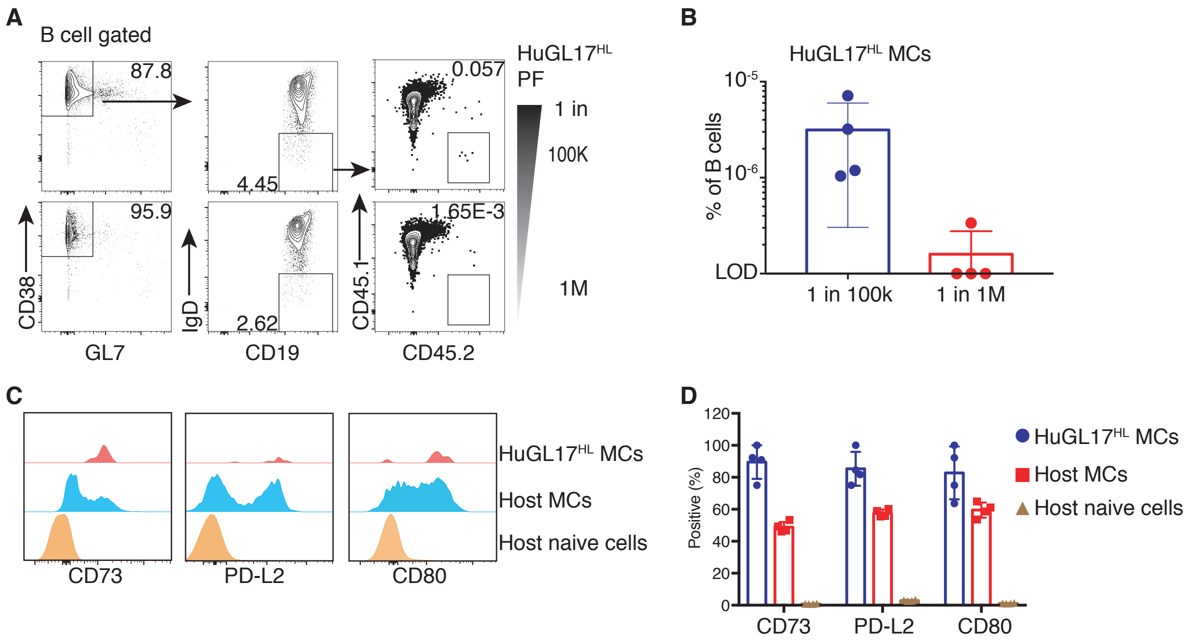
A
B cell gated
87.8
95.9
4.45
2.62
0.057
1.65E-3
HuGL17HL PF
1 in
100K
1M
CD38
GL7
IgD
CD19
CD45.1
CD45.2
B
HuGL17HL MCs
% of B cells
10-5
10-6
LOD
1 in 100k
1 in 1M
C
HuGL17HL MCs
Host MCs
Host naive cells
CD73
PD-L2
CD80
D
Positive (%)
120
100
80
60
40
20
0
CD73
PD-L2
CD80
HuGL17HL MCs
Host MCs
Host naive cells

Figure S4

**Fig S4**. HuGL17 B cells were seeded at 1 in 1 million or 1 in 100,000, challenged with eOD-GT8, and HuGL17 CD45.2 memory B cells were analyzed at d49. **(A)** Gating strategy to enumerate $CD45.2^{+}CD19^{+}CD38^{+}IgD^{-}GL7^{-}$ ("memory B cells"). **(B)** Enumeration of memory B cells as a percentage of total B cells. Each datapoint represents the value obtain in one mouse. **(C,D)** Evaluation of memory B cell expression of CD73, PD-L2 and CD80 showing that HuGL17 cells were largely triple positive.

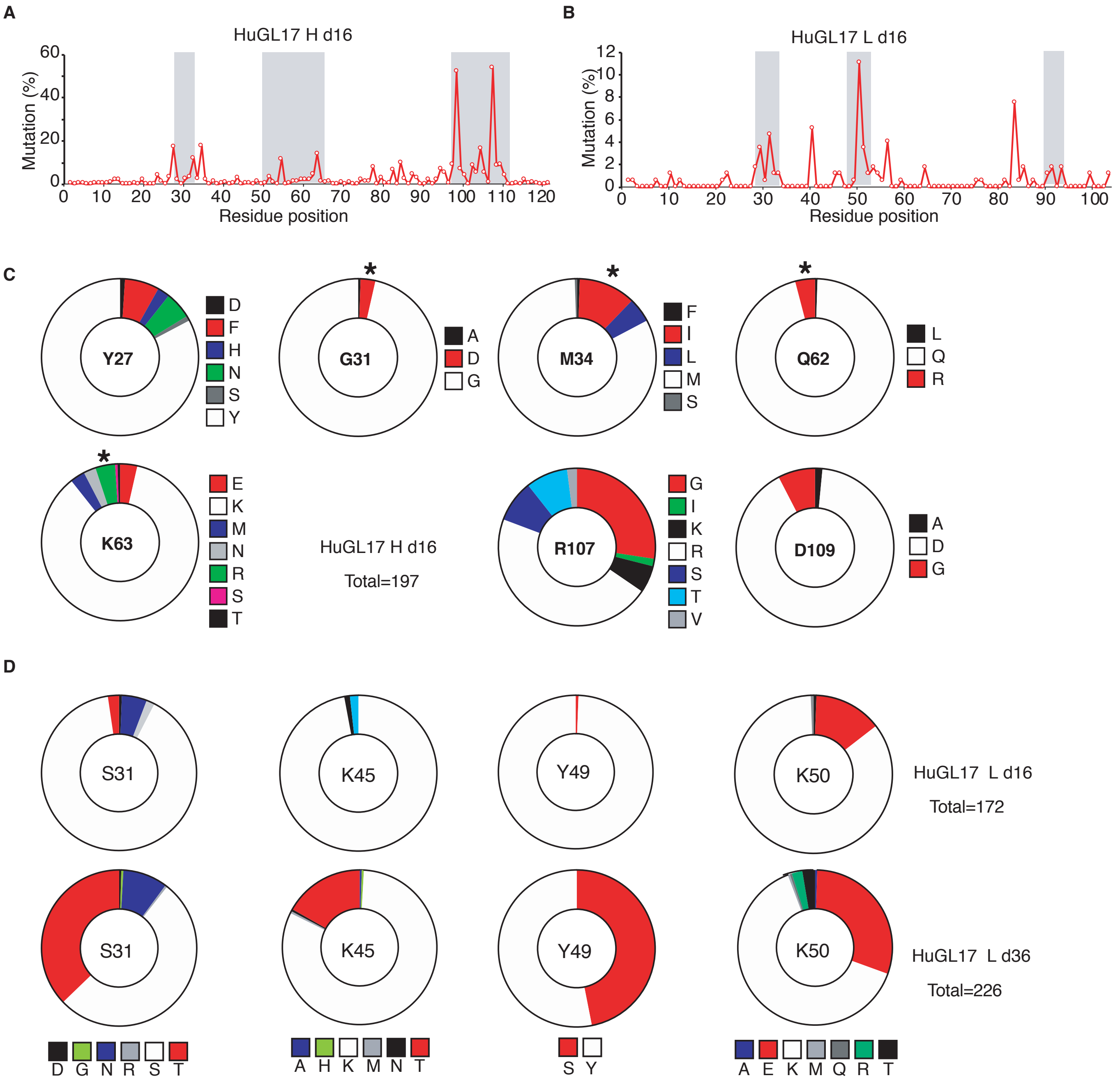
A
HuGL17 H d16
Mutation (%)
Residue position
B
HuGL17 L d16
Mutation (%)
Residue position
C
Y27
G31
M34
Q62
K63
R107
D109
HuGL17 H d16
Total=197
D
S31
K45
Y49
K50
HuGL17 L d16
Total=172
HuGL17 L d36
Total=226

Figure S5

**Fig S5**. Further analysis of somatic mutations of HuGL17 cells. **(A,B)** Analysis of the extent of mutation on HC and LC on d16 as a function of amino acid position. **(C)** Prominent HC mutations at d16. Asterisks indicate VRC01-class mutations. The extent and variety of changes at R107 suggested selection against R at this position. **(D)** Comparison of prominent LC mutations at d16 and d36. Total sequences analyzed are indicated.

**Table S1.** Features of VRC01-class germline BCRs identified in human PBMC compared to a VRC01 iGL.

| Name | H-chain V/D/J | | | CDRH3 | L-chain V/J | | CDRL3 | GT8 $K_D$ |
|---|---|---|---|---|---|---|---|---|
| HuGL16 | VH1-2*02 | D6-13*01 | JH4*02 | ARVRYGSWTGYYFDY | VK1-33*01 | JK4*01 | CQQYDLF | 18.5 μM |
| HuGL17 | VH1-2*02 | D6-13*01 | JH4*02 | ARVIRSSSSWRYDY | VK1-05*03 | JK1*01 | CQQYETF | 1.3 μM |
| HuGL18 | VH1-2*02 | D6-13*01 | JH4*02 | ARDHQGHSSSWSKRFDY | VK3-20*01 | JK1*01 | CQQYETF | 125 nM |
| gl-VRC01 | VH1-2*02 | D2 | JH1*01 | ARGKNSDYNWDFQH | VK3-15*01 | JK2*01 | CQQYEFF | 30 pM |
